# Multi-physics modeling for ion homeostasis in multi-compartment plant cells using an energy function

**DOI:** 10.1101/2025.02.11.637607

**Authors:** Guillaume Mestdagh, Alexis De Angeli, Christophe Godin

## Abstract

Plant cells control their volume by regulating the osmotic potential of their cytoplasm and vacuole. Water is attracted into the cell as the result of a cascade of solute exchanges between the cell subcompartments and the cell surroundings, which are governed by chemical, electrostatic and mechanical forces. Due to this multi-physics aspect and to couplings between changes of volumes and chemical effects, modeling these exchanges remains a challenge that has only been partially adressed. In this paper, we introduce an energy-based approach to couple chemical, electrical and mechanical processes taking place between several subcompartments of a plant cell. The contributions of all physical effects are gathered in an energy function that allows us to derive the equations satisfied by each variable in a systematical way. This results in a modular, unified approach that can be interpreted analytically. We illustrate these properties on the modeling of ion and water transport in a guard cell during stoma opening. We represent the stoma opening process as a quasi-static evolution driven by hydrogen pumps in the plasma and vacuolar membranes, resulting in an interpretable model with few parameters. We perform numerical simulations to investigate the role of each hydrogen pump in this process. We show that this energy-based approach allows us to highlight a hierarchy between the forces involved in the system, that can be exploited to interpret the emergent properties of this complex multi-physics system.

## 1 Introduction

Plant cells change their size and shape to achieve various key living functions. These changes can be irreversible, as in growing cells whose increase in volume triggers wall remodeling and cell division. They can also be reversible, as in the case of guard cells. Located on the surface of plant leaves and stems, guard cells work in pairs to control the aperture of small pores called stomata. These specialized cells are able to repeatedly get inflated and de-inflated to open and close stomata, and regulate plant transpiration and carbon dioxide absorption (Roelfsema and Hedrich, 2005; Eisenach and De Angeli, 2017).

Like most plant cells, guard cells control their volume by regulating the amount of water entering their vacuole (Martinoia et al., 2012; Krüger and Schumacher, 2018; Kaiser and Scheuring, 2020; Dünser et al., 2022). For this, they actively control the osmotic potential of their cytoplasm and vacuole by importing or exporting ions across the membranes delimiting these compartments (the plasma and vacuolar membranes, respectively). These ion transfers must be precisely synchronized over short time scales to achieve the inflation or de-inflation of the cell. Ions are transported by a wide variety of specialized proteins located on each membrane, including pumps, channels, exchangers, etc. Due to their specific stoichiometry, these transport mechanisms define constraints on possible transfers of solute between subcellular compartments. The forces that drive ion transfer are of multiple natures. They integrate chemical, electrical, and mechanical components, whose combination determines the dynamics of this multi-physics, multi-membrane system.

Modeling approaches have been widely used to help analyze the behavior of such complex systems, as a complement to experimentation. A first category of approaches focuses on electrochemical exchanges across membranes using ordinary differential equations. In these models, the variation of ion concentrations in each cell compartment is defined by the ion fluxes crossing membranes through a range of transport proteins. Vitali et al. (2016), for example, identify the most significant parameters controlling the volume dynamics of beetroot vacuoles, and they use simulation to explore the effect on the vacuole volume of a range of values for these parameters. In a recent work, Li et al. (2024) account for the multimembrane structure of the cell, and simulate the ion fluxes across both the plasma and vacuole membranes of a guard cell in response to variations in the external potassium and chloride concentration. Note that similar studies have also been carried out to model ion transport in animal or fungi cells (Kahm et al., 2012; Astaburuaga et al., 2019). Many cell physiology models use a formalism inspired by chemical kinetics to represent the dynamic of each transport process. Gerber et al. (2016) opt for a different formalism inspired by thermodynamics, introduced in the 1950s (Staverman, 1951, 1952; Kedem and Katchalsky, 1958), where the flux across a membrane is proportional to the electrochemical potential gradient between the two connected compartments. They use a parameter estimation procedure to fit their simulation results onto experimental measures. It is interesting to note that the variety of modeling approaches used to determine the flux of ions between compartments illustrates the fact that the modeling of transport dynamics in cells remains a challenging and discussed topic.

A second category of approaches aims at modelling the mechanical response of the cell wall to an increasing turgor pressure inside the cell. In the case of guard cells, many studies examine the cell wall elastic properties, with a combination of modeling and experiments (see the review by Woolfenden et al., 2018). Meckel et al. (2007) for instance discuss whether guard cells widen or. elongate as their volume increases. By comparing finite element simulations and experimental measures, Woolfenden et al. (2017) estimate the stiffness parameter distribution of the cell wall. Similar approaches are followed by Marom et al. (2017) and Yi et al. (2018). Carter et al. (2017) study the role of wall stiffening at the extremities of guard cells in the stoma opening process, while Jaafar et al. (2024) focus on the role of cell wall anisotropy.

The models mentioned above either describe electrochemical or mechanical phenomena. On the one hand, ion exchange models involve an osmotic pressure but rarely describe water movements between compartments, whereas on the other hand, mechanical models predict cell deformation as a result of the turgor pressure whose origin is not specified. The interplay between these two categories of effects is often not taken into account, as it represents an additional difficulty. Also, the multi-compartment aspect of plant cells is rarely considered. Recent studies by Cubero-Font and De Angeli (2021) and Li et al. (2024) highlight the need to include it in modeling approaches, to better understand how cells synchronize the regulation of their membranes. This new trend in plant cell physiology calls for models and formalisms able to provide novel insight into multi-compartment systems.

The challenge of building multi-membrane models that couple electrochemistry with mechanics is addressed in the OnGuard model (Hills et al., 2012). This well-established framework aims to provide realistic simulations of solute exchanges in a guard cell. To connect the guard cell physiology with the stoma aperture, the authors use a linear pressure-volume law, along with a linear volume-aperture relation, whose coefficients are empirically set based on experimental results. OnGuard has been implemented as a software platform, that has proven useful to reproduce experimental results (Chen et al., 2012), and to drive new experiments by predicting guard cell behavior (Wang et al., 2012; Minguet-Parramona et al., 2015; Vialet-Chabrand et al., 2017). To run a simulation, the user chooses among an exhaustive list of membrane transport proteins and sets the dynamic parameters of each reaction law. Results are available through different plots showing the time evolution of the guard cell volume and aperture, osmolyte concentrations and fluxes, among many other physical quantities.

In this paper, similarly to OnGuard, our main objective is to develop a multi-membrane, multi-physics model for the regulation of plant cell volume control. However, we follow a different and complementary approach. While OnGuard focuses on exhaustiveness and resemblance to experimental results, we center our attention on interpretability and modularity. We seek to integrate electrochemical and hydro-mechanical processes in an explicit and mechanistic way, while keeping the model interpretable. To meet these ambitions, we unify the various physical processes by gathering their contributions into a common global energy function. This energy function naturally brings some modularity to the model, as its components can be added or removed without impact on the other components of the system. In addition, it keeps generic features (e.g. implementation of the main physical processes) well separated from specific ones (e.g. choice of specific transport stoichiometry, etc.). We exploit this approach to simulate ion transfer between the subcellular compartments of a guard cell during stoma opening. To keep the model simple, we assume that the underlying physical processes follow at every moment a moving equilibrium, which progressively evolves in response to successive perturbations. This makes it possible to neglect the dynamics of ion transport and to keep fewer parameters in the model. In such quasi-static simulations, time-dependent ordinary differential equations are replaced with a series of nonlinear equations that are used to compute the sequence of equilibrium states. Altogether, the use of an energy landscape and quasi-static simulation allows to obtain a simple geometrical interpretation of the system evolution. We will show that the resulting formulation is flexible and easily provides physical insight into the coupling of multiple physical processes by giving access to the role of the different parameters in the system’s regulation.

## 2 A multi-physics model based on an energy function

We now introduce the formalism used to construct a simple representation of a plant cell. We reduce the cell to a set of nested compartments connected by transporters. For clarity reasons, we use the term *transporter* to designate any process that transports reactants across a membrane. This definition includes pumps, channel, coupled ion transport, and even water permeating across membranes. As mentioned in Section 1, we seek to compute a sequence of equilibrium states for the system, that is parametrized by the chemical extent of some controlled transport reactions. In other words, a resulting equilibrium state for the whole system is associated with a given state of the active transport processes. As active transporters work, the system equilibrium state constantly adjusts in consequence.

From now on, we reduce an object as complex as a plant cell to a *multi-membrane complex*, which is an arrangement of one, two, or possibly more nested compartments enclosed by membranes. Each compartment contains a certain chemical amount (in mol) of chemical species such as ions and water. In this article, water can circulate across membranes, while the transport of ions between compartments is allowed by several active and passive transporters. The multi-membrane complex is surrounded by an external environment with fixed ion concentrations, which acts as a reservoir. A plant cell is typically represented by a complex with two compartments (the cytoplasm and the vacuole). For instance, the system shown in Figure 1, involves two membranes, four chemical species (hydrogen, chloride, potassium, water) and eight transporters.

**Figure 1:**
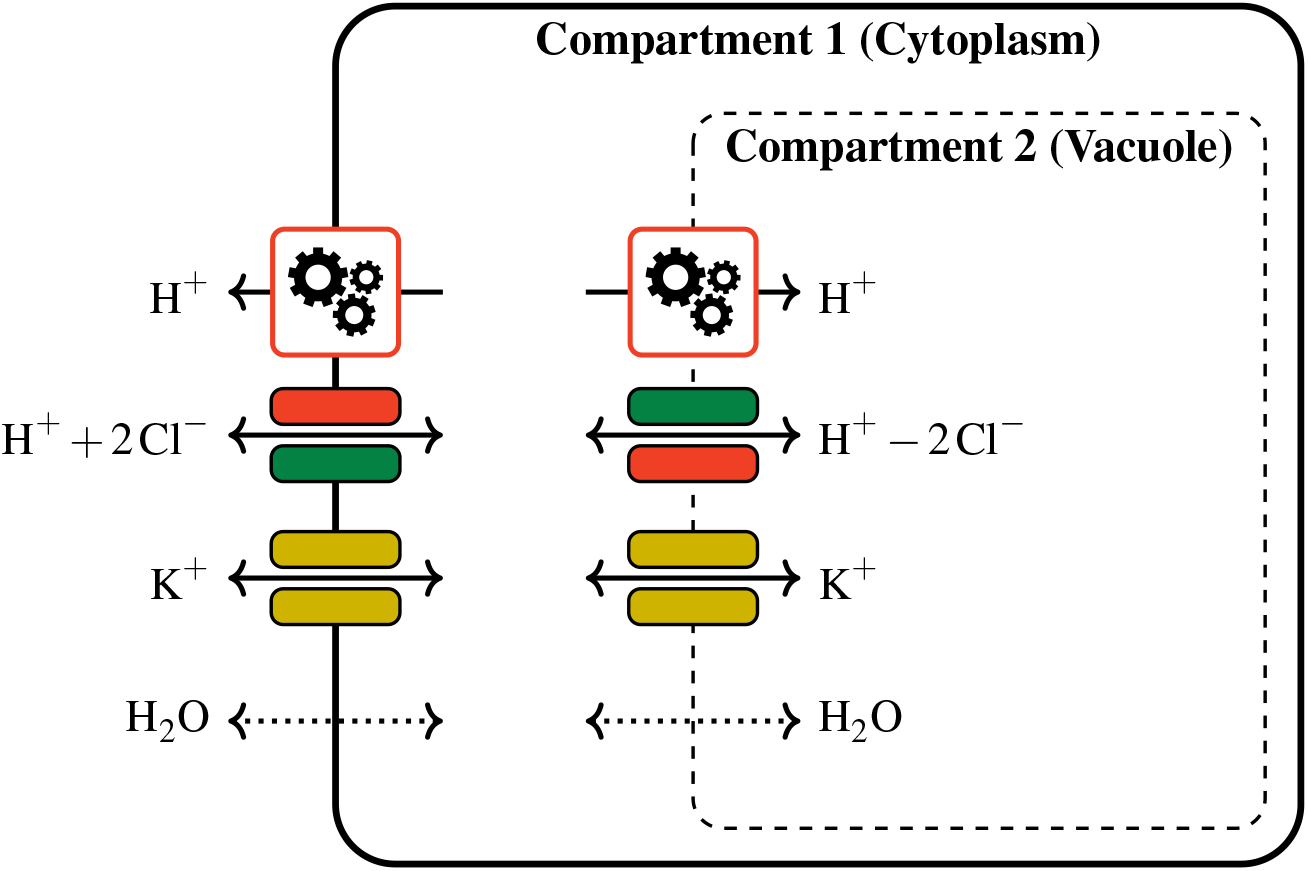
Representation of a guard cell as a multi-membrane complex. The model involves two membranes and a hand-picked selection of transporters. Each compartment contains a certain amount of hydrogen, chloride, potassium and water. All these chemical amounts compose the system state. Mechanical energy can only be stored into the outer membrane, which results in an equal pressure in both compartments.

The state of the system at a given moment is entirely defined by the chemical amount of each reactant (i.e. each considered chemical species) in each compartment. Of course, to perform simulations involving a multi-membrane model, some fixed parameters need to be defined, such as the initial content of all compartments or initial electric potentials across membranes, but these parameters are not variables, and they remain fixed along the simulation. The chemical amount of reactant *A* in compartment *i* is denoted by *n*_*i*,*A*_, and all chemical amounts are gathered in the state vector **n** ∈ ℝ^*N*^. Here, the size *N* of the state vector is the product of the number of reactants with the number of compartments. For instance, the state vector for the model from Figure 1, of size *N* = 8, reads

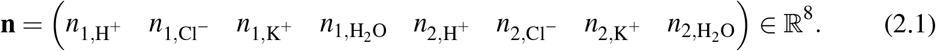

Note that the variables in the state vector are first ordered by compartment, and then by reactant.

An equilibrium state is defined as the state that minimizes an energy function among a range of states attainable by the system. More formally, the equilibrium state is the solution to a constrained optimization problem of the form

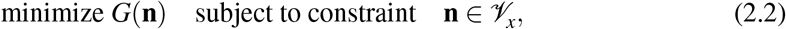

where *G* is an energy function to minimize, **n** is the variable that describes the system state, and *V*_*x*_ is the range of states attainable by the system. Below, we give details about the various components of this minimization problem, starting with the energy function.

### 2.1 Energy function and potentials

To understand the dynamics of a physical system is to understand how its protagonists store, exchange or dissipate energy. In particular, the energy that is stored and does not correspond to a current motion is called potential energy. It is the energy paid by the system just to be in a certain state. The potential energy function of a system depends on the system state only, and it describes the forces that influence the system. These forces make the physical system spontaneously evolve to decrease its potential energy, generally transforming it into heat. A state where the potential energy is at a local minimum is called an equilibrium state. When a physical system has reached an equilibrium, it ceases to evolve, and when it is taken away from this equilibrium, it tends to go back to this equilibrium (we will only meet stable equilibria in this study). From now on, we simply use *energy function* to refer to the potential energy function. The potential energy function should not be mixed up with potentials (chemical potential, electric potential), that rather correspond to derivatives of the energy function. Namely, a chemical potential is expressed in J*/*mol, an electric potential in J*/*C and a pressure in J*/*m^3^, but the potential energy is expressed in J.

Defining an energy function for a physical system is a very convenient modeling practice, as energy is a common currency between a wide range of physical processes. In general, the system energy function is the sum of several terms, with each term corresponding to a specific physical process. In this paper, the energy function, is split into three terms, as we consider three types of physical effects. First, chemical effects tend to equilibrate the concentration of reactants between both side of each membrane. Second, electrostatic effects oppose to the building of large electric potential variations across membranes. Finally, mechanical effects act on volume changes.

Though the energetic formalism is compatible with several choices of shape for the cell, geometry plays a significant role when it comes to deriving an actual expression for energy function terms. In this paper, we stick to a very simple geometry for each membrane, namely a cylinder that elongates in one direction only, along its revolution axis, so that one-dimensional considerations are sufficient to define the elastic deformation energy. When the cell volume changes, only the cylinder lateral surface is subject to strain in the direction of the elongation, and the cylinder volume and lateral surface area remain proportional to its length. We denote by *V*_*i*_(**n**) the volume enclosed in the *i*-th membrane, and by *A*_*i*_(**n**) the area of its lateral surface. For a cylinder with fixed radius *r*, the lateral surface area and the volume satisfy *A*_*i*_(**n**) = 2*V*_*i*_(**n**)*/r*. Note that the *i*-th membrane encloses compartment *i*, but also all compartments with index *j > i*.

We now give a little more details about each term of the energy function.

#### Chemical energy

The first term we consider measures a discrepancy between the system state and a chemical equilibrium. Chemical forces usually determine in which direction a chemical reaction occurs (Nelson, 2004, Chapter 8). Though we do not consider any chemical transformation in this model, chemical forces still play a role in determining which way ions circulate through a transporter. Namely, the chemical energy takes larger values when the concentration of a reactant is very different between two compartments, and chemical forces tend to promote, for each reactant, a uniform concentration among compartments. Chemical forces are responsible for the osmosis process, i.e. attracting water in compartments where the solute concentration is high.

We use the standard expression for the chemical energy term, given by

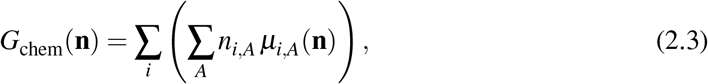

where *µ*_*i*,*A*_(**n**) is the chemical potential associated with reactant *A* in compartment *i*. When chemical transformations occur, the chemical potential determines how the chemical energy changes when reactant *A* appears or disappears out of the blue in compartment *i*, and its classical expression reads

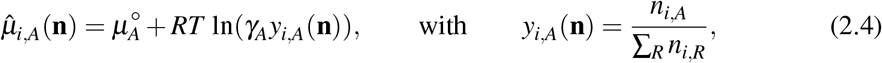

where *y*_*i*,*A*_(**n**) the mole fraction of reactant *A* in compartment *i*, and 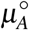 and *γ*_*A*_ represent the chemical potential of pure species and the activity coefficient of reactant *A*, respectively (Mortimer, 2008, Section 6.3). However, we only consider solute *transport* in this study, and an increase of *n*_*i*,*A*_ without variation of other quantities means that some reactant *A* has been taken from the external environment and transported into compartment *i*. The corresponding marginal variation rate of chemical energy, denoted by

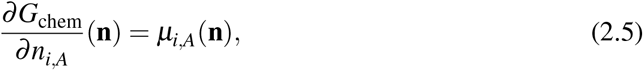

involves the chemical potential of *A* in the external environment. It reads

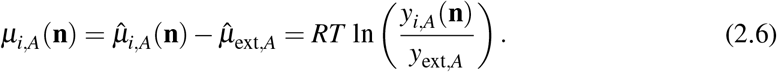

Here, the external environment is considered as a reservoir, where each reactant *A* is present with the constant mole fraction *y*_ext,*A*_, so that the chemical potential 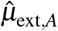 of *A* in the external environment is constant. Also, note that *µ*_*i*,*A*_ only depends on chemical amounts in compartment *i*, so that (2.3) is actually made of independent terms, which each term depending on mole fractions in one compartment. From expressions (2.3) and (2.6), we can check that the chemical energy function is at a minimum when all chemical potentials are zero, i.e. mole fractions in each compartment are equal to external mole fractions.

#### Electrostatic energy

The electrostatic term of the energy function aims to penalize large differences of electric potential across each membrane. When a charged particle crosses a membrane from one compartment to another, the net transfer of electric charge results in a charge imbalance across the membrane and a difference of electric potential between the two compartments. This phenomenon, also known as membrane polarization, tends to oppose to the passage of more similar charges in the same direction. From a modelling point of view, a membrane can be considered as an electric capacitor, i.e. a thin insulating layer between two charged compartments, allowing charges to accumulate along the membrane when a difference of potentials occurs. As a consequence, the electrostatic energy value is the sum of the energy of all capacitors. The electrical capacitance of a membrane is proportional to its area. The capacitance per unit membrane area in biological cells is typically 1 *µ*F*/*cm^2^ (Keener and Sneyd, 2009, Section 2.6).

The electric structure of a multi-membrane complex can be represented by an equivalent circuit, shown in Figure 2 in the two-membrane case. Each compartment corresponds to a node in the circuit, while the external environment is represented by the ground. Compartments are connected by capacitors that represent the membranes, while the two current sources symbolize the action of adding charges from the external environment into one of the compartments. To understand the relationship between the content of each compartment and the state of capacitors, let us apply Kirchhoff’s circuit laws: if we add a charge d*q*_1_ into compartment 1 and d*q*_2_ into compartment 2, then the charge stored in capacitor *C*_2_ changes by d*q*_2_, while the charge stored in capacitor *C*_1_ changes by d*q*_1_ +d*q*_2_ (here, charge displacements act like current intensities). In other words, the charge *Q*_*i*_(**n**) stored in the capacitor *C*_*i*_ depends on the cumulated electric charge in all compartments enclosed inside the *i*-th membrane. In particular, transporting charged particles from compartment 1 to compartment 2 corresponds to the situation d*q*_1_ = −d*q*_2_, which does not change the charge stored in capacitor *C*_1_.

**Figure 2:**
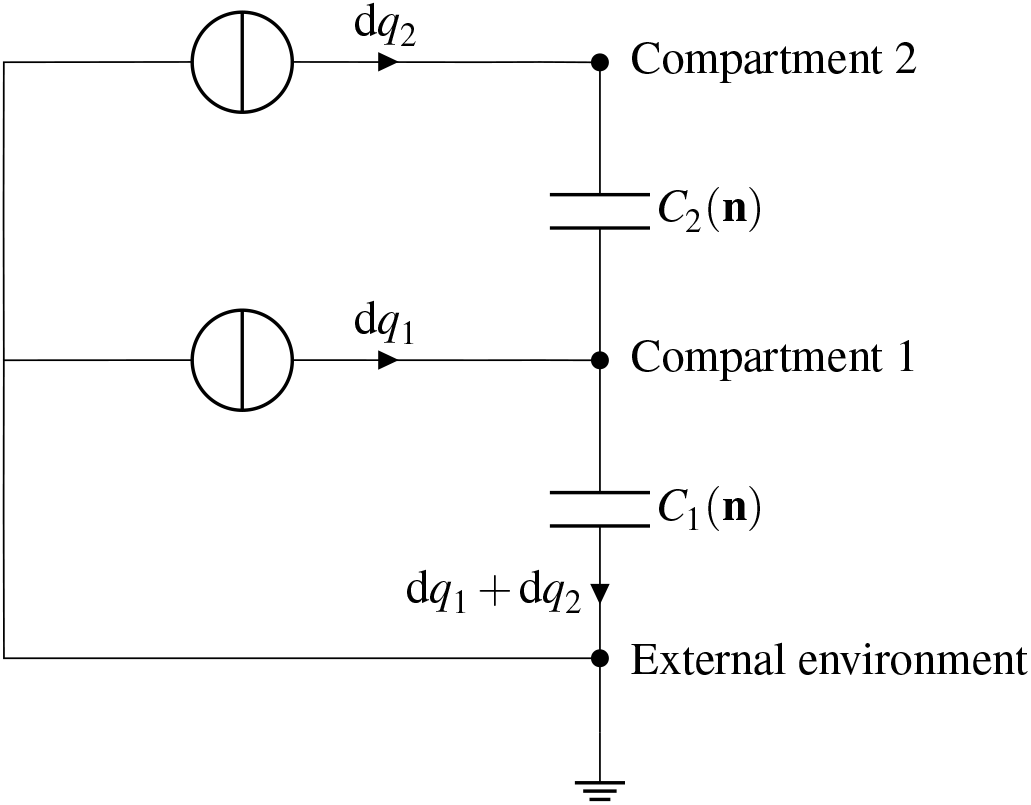
Electric representation of a two-membrane complex, where each capacitor represents a membrane and each compartment corresponds to a node in the circuit. Variations of ion amounts in each compartment result in adding or removing charges in these compartments. The circuit representation suggests that adding charged particles into compartment 2 contributes to charging both capacitors *C*_1_ and *C*_2_.

Let us now derive an expression for the electrostatic energy stored in capacitor *C*_*i*_. If we denote by **n**_*k*_ = (*n*_*k*,*A*_, *n*_*k*,*B*_, …) the vector of all chemical amounts in compartment *k*, the charge contained in compartment *k* is defined by

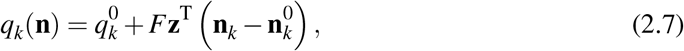

where **z** = (*z*_*A*_, *z*_*B*_, …) stores the charge of each reactant, and *F* is the Faraday constant. The parameters 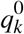 and **n**^0^ are fixed as part of the initial conditions. The expression for the charge stored in capacitor *C*_*i*_ reads

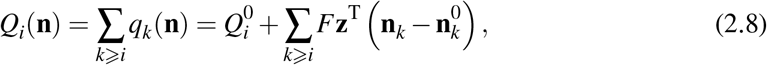

where 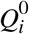 denotes the stored charge in initial conditions. The energy stored in capacitor *C*_*i*_ is defined using the classic formula

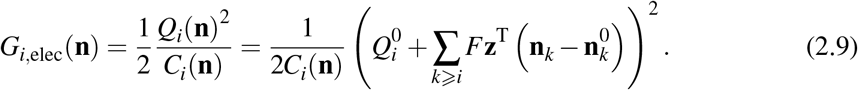

Note that the initial membrane charge 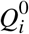 can be deduced from experimental measures of initial membrane potentials. The expression for the capacitance *C*_*i*_(**n**) depends on the cell geometry. As mentioned above, the cell is modelled as a cylinder, and for simplicity we only consider its lateral surface to evaluate the membrane capacitance, which is defined by

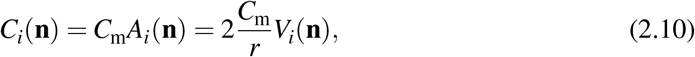

where *C*_m_ is the capacitance per unit surface.

Finally, we evaluate the total electrostatic energy by adding up the energy stored in each capacitor, i.e.

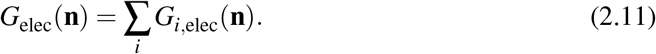

#### Mechanical energy and cell geometry

The mechanical energy term represents the elastic energy stored in the cell wall as the cell volume increases. While cell growth is an irreversible process involving cell wall elasticity and plasticity at the same time, guard cell deformation during stoma opening is reversible, as the stoma opening/closing cycle happens several times, at least once a day. As we focus in this paper on the quasi-static modelling of a guard cell, the only mechanical effect we consider is elasticity.

The mechanical energy stored in a membrane is a function of the volume it encloses. When subject to a volume change, the membrane adopts a resulting mechanical configuration, which corresponds to a value for its mechanical energy. In this paper, the only contributor to compartment volume is water, so that the volume enclosed in the *i*-th membrane is defined by

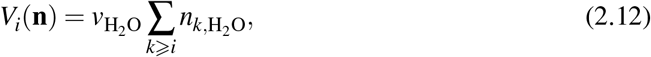

where 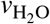 is the molar volume of water. Keep in mind that *V*_*i*_(**n**) is not the volume of compartment *i*, but the cumulated volume of all compartments enclosed in the *i*-th membrane. Also, in the context of plant cells, we assume that the only membrane that offers mechanical resistance to volume changes is the outer membrane (membrane 1), as in plant cells the plasma membrane is coupled with the cell wall, while the vacuole membrane (membrane 2) is only a lipid bilayer. The resulting elastic deformation energy only depends on the whole cell volume *V*_1_(**n**), which is the sum of all compartment volumes. In other words, defining the mechanical energy term boils down to defining a function of one variable, sometimes referred to as a pressure/volume law. In terms of pressure, it means that no membrane except the first one applies mechanical forces on the fluid it contains, and the mechanical pressure is the same in every compartment. If we define the pressure *P* = *∂G*_mech_*/∂V*_1_, by looking at (2.12) we obtain

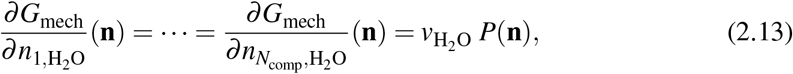

which means that it has the same cost in mechanical energy to add some water into any compartment.

Without choosing a cell geometry, it is difficult to tell much more about the mechanical energy term. We now exploit the cylindrical cell shape to come up with an expression. When the cell expands, the strain in the lateral surface is given by the expression *ε*(**n**) = *V*_1_(**n**)*/V*_1_(**n**^0^) − 1. We denote by Σ^0^ the initial cylinder lateral surface and by *A*_*i*_(**n**^0^) its area. Using a linear constitutive law, the energy integrated over the whole lateral surface reads

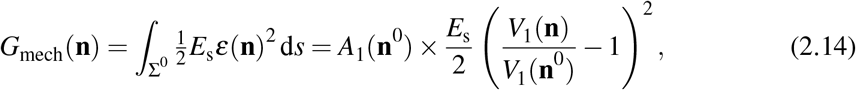

where *E*_s_ is a surface elastic modulus (in Pa · m).

Though we used a simple one-dimensional model for the cell geometry, the energetic formalism is completely compatible with finer geometries, including alternative descriptions of the cell expansion (Meckel et al., 2007) or even finite-element based geometries (Woolfenden et al., 2017, 2018). In this last case, an explicit formula for the mechanical energy term might not be available.

#### Enforcing equilibrium in starting conditions

To improve consistency between the model and experimental conditions, we need to start a simulation from any user-specified initial state **n**_0_, which can be a state measured at the beginning of an experiment. However, in a quasi-static model, **n**_0_ must be an equilibrium state of the system, i.e. it should correspond to a minimum of the energy function *G*. To make sure that **n**_0_ represents an equilibrium state, we add a linear correction term 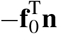 to the energy function *G*, where **f**_0_ is adjusted so that ∇*G*(**n**_0_) = 0, which means that **n**_0_ is a minimum of *G*. Namely,

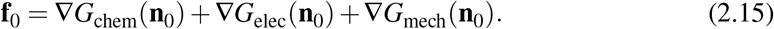

The force **f**_0_ can be interpreted as the contribution of the environment to the energy function. The guard cell is the theater of many processes, associated to metabolism or other functions that are not explicitly modelled. From the point of view of our model, we assume that all this activity can be summarized by the constant force **f**_0_. For a given **n**_0_, the total energy function that is actually minimized during the simulation is defined by

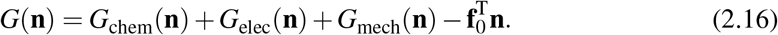

Though this energy function depends on **n**_0_, we keep calling it *G* in the remaining of this section, as **n**_0_ is a fixed parameter of the model.

### 2.2 Membranes, transporters, directions

The energy function described in Section 2.1 defines the forces that influence the system. It is completely agnostic of the disposition of transporters. If the system was free to reach any state **n** ∈ ℝ^*N*^, it would converge toward the global minimum of the energy function *G*. However, some states cannot be reached by the system, as chemicals can only circulate between compartments through transporters, which exhibit a specific stoichiometry. The disposition of transporters defines the set of states attainable by the system, denoted by *V*_**x**_ in (2.2).

In this subsection, we describe the connection between the disposition of transporters and the set of attainable solutions. For the sake of simplicity, we use a simple one-membrane complex to illustrate the chosen formalism. This toy system, shown in Figure 3, involves one compartment, three chemical species (hydrogen, chloride, water) and three transporters. The same formalism will later be extended, *mutatis mutandis*, to a complex with several membranes.

**Figure 3:**
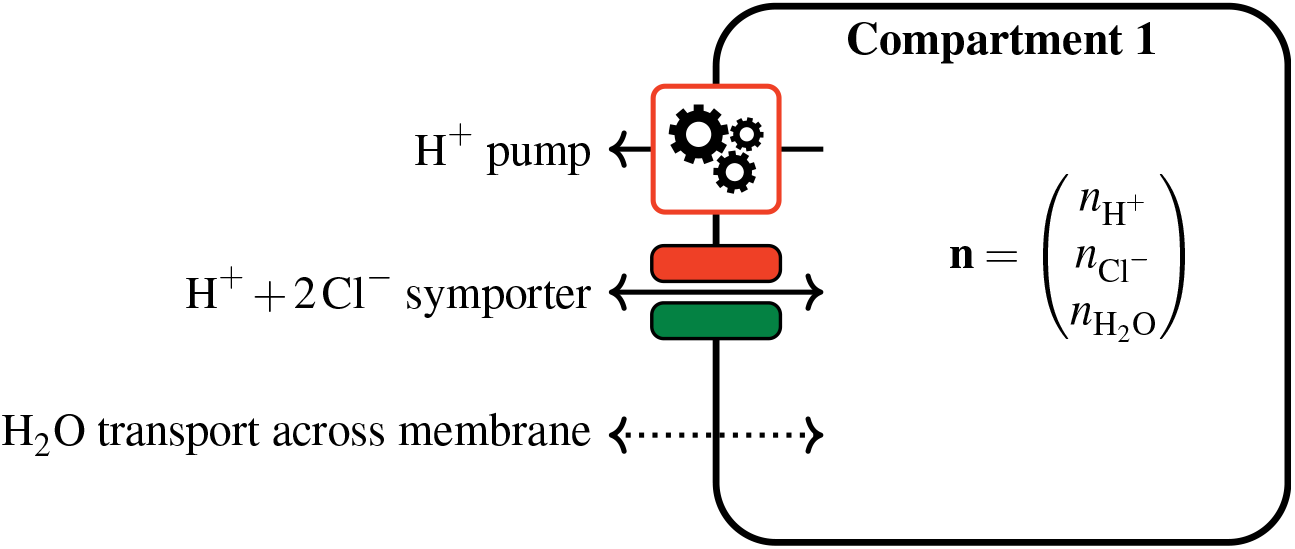
A toy multi-membrane complex with one membrane and three transporters. The hydrogen pump is an active transporter, while the hydrogen/chloride symporter is passive. Water transport across the membrane is also considered passive. As there are three species and one compartment, the system state is a vector of size 3.

#### Points and directions in the state space

In the one-membrane complex, the system state is defined by the chemical amount of each species *A* in the single compartment. We use the slight abuse of notation *n*_*A*_ = *n*_1,*A*_, so that the state vector now reads

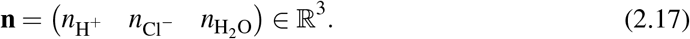

The vector **n** may be interpreted as a point in a three-dimensional state space, whose coordinates are the amounts of reactants. Also, a variation of reactant quantities corresponds to a displacement along a certain direction in this space, represented by a vector of size 3. For instance, in Figure 3, the first transporter is a hydrogen pump. It ejects hydrogen from the compartment to the external environment, which results in the hydrogen amount 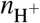 decreasing. As 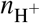 is the first coordinate of the state vector **n**, this is represented in the state space by **n** evolving along the vector

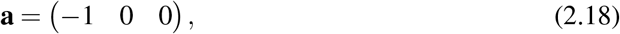

where we have used the same ordering as (2.17) for chemical species. Once the hydrogen pump has ejected *x* moles of hydrogen (and no other transport occurred), the new state now reads

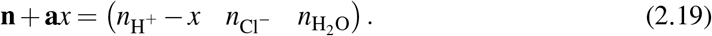

The second transporter is called a symporter because it transports two species in the same direction, at the rate of one hydrogen ion for two chloride ions, while the third transporter symbolizes water permeating across the membrane. In a similar fashion as previously, each one of these two transporters allows the state to evolve in a particular direction. These directions, denoted by **v**_1_ and **v**_2_, are defined after the stoichiometry of the two transporters (see Figure 3), namely

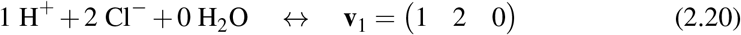

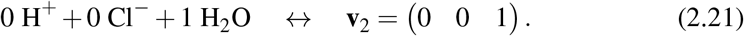

In a general multi-membrane complex, the state can only evolve in directions defined by the stoichiometric coefficients of transporters, which restricts the range of attainable states. For instance, in the toy model, if we block the hydrogen pump, then the amount of chloride cannot change independently of hydrogen, as the only way out for chloride is through the hydrogen/chloride transporter. In particular, the value 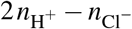 must remain constant. Here, the state **n** can only move along two directions **v**_1_ and **v**_2_, and therefore the set of attainable states is a subspace of dimension two.

#### Constrained energy minimization

The hydrogen pump is a particular transporter: whereas the other transporters just passively allow ions and water to cross the membrane at the whim of electrochemical potentials, the pump consumes some energy from an external source to eject hydrogen from the compartment, sometimes against chemical potentials. We call the hydrogen pump an *active* transporter while other transporters are *passive*. This distinction is critical for the remaining of this section, as the directions associated with passive and active transporters play different roles in the definition of *V*_**x**_. The stoichiometry of passive transporters defines the directions that span *V*_**x**_, while active transporters define its displacement in directions orthogonal to *V*_**x**_.

Let us assume for a moment that the active pump is not working, which means that hydrogen and chloride can only circulate together through the hydrogen/chloride transporter. Starting from an initial state **n**^0^, the system evolves toward states of lower energy, but, as mentioned above, the state vector **n** can only move along the two linearly independent directions **v**_1_ and **v**_2_. In other words, **n** is restricted to an affine plane *V*_0_ ⊂ ℝ^3^ containing **n**^0^. If we denote by **w**_1_ = **v**_1_ *×* **v**_2_ = (2, −1, 0) a normal vector to *V*_0_, the subspace of attainable states is characterized by the equation

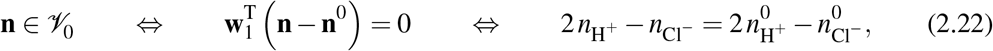

where 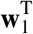 is the transpose of **w**_1_. As a consequence, when the active transporter does not work, the equilibrium attained by the system is the minimizer of the energy function *G* among the subspace *V*_0_. Now, assume that the hydrogen pump ejects *x* moles of hydrogen before stopping, bringing the system to the state **n**^0^ + **a***x* (see 2.19). To find an equilibrium, the state **n** is now restricted to another subspace, denoted by *V*_*x*_,whose equation reads

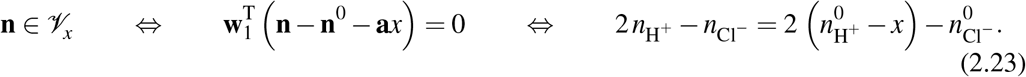

It is notable that, for *x* ≠ *x′*, the subspaces *V*_*x*_ and 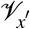 are disjoint (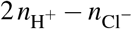 takes different values), and as a consequence, the resulting system equilibria are distinct.

Figure 4 shows a slice view of the space of states in the 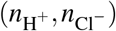 plane, with representations of the subspaces *V*_*x*_ corresponding to several values of *x*. The lines in the background represent the level curves of the energy function *G*. The point that minimizes *G* in a given sub-space *V*_*x*_ is the point where a level curve is tangent to *V*_*x*_. As *V*_*x*_ move, the equilibrium state **n**^∗^(*x*) is pushed toward higher levels of energy, following the red curved trajectory.

**Figure 4:**
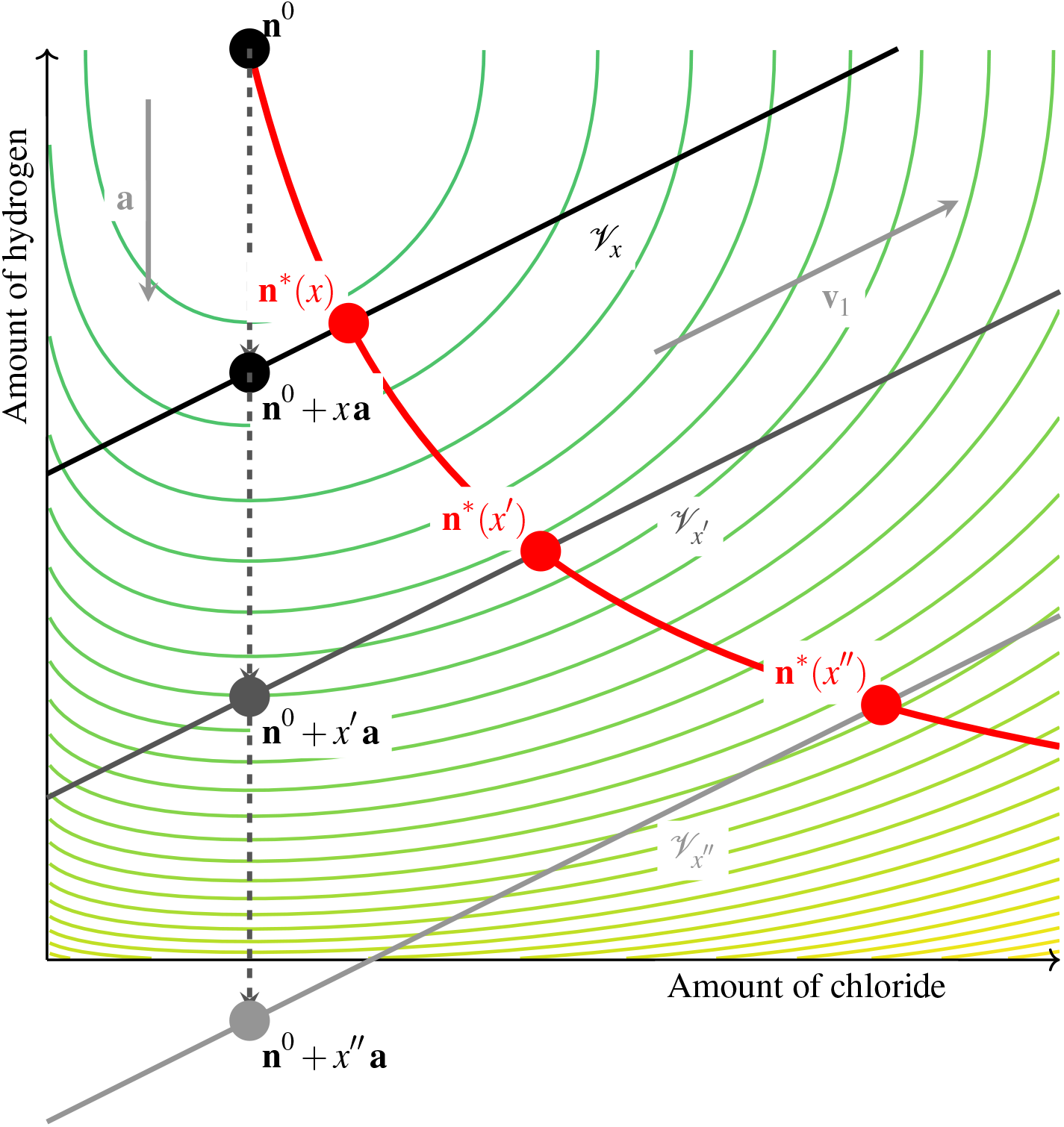
Slice view of the space of states in the plane 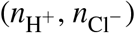 for the toy problem. The level curves in the background represent the energy function *G*. Once the hydrogen pump has brought the system from **n**_0_ to **n**_0_ + *x* **a**, the passive evolution toward an equilibrium is only possible along the affine subspace *V*_*x*_, which is directed by **v**_1_ and **v**_2_ (**v**_2_ is out of the plane). As *x* increases, the plane *V*_*x*_ continuously moves, dragging the equilibrium state **n**^∗^(*x*), which follows the thick curved trajectory.

In the toy model from Figure 3, the extent of active transport *x* is the input variable, while the output is the corresponding equilibrium state **n**^∗^(*x*). When the amount of hydrogen ejected by the pump describes an interval [0, *x*], it progressively displaces the passively attainable subspace *V*_*x*_, and the system equilibrium is continuously brought from **n**^∗^(0) ∈ *V*_0_ to **n**^∗^(*x*) ∈ *V*_*x*_. A more general multi-membrane complex might feature several active transporters, whose chemical extents are stored in a vector **x** = (*x*_1_, *x*_2_, …), and whose corresponding directions are stored as the columns of a matrix **A** = (**a**_1_|**a**_2_| · · ·). In that case, the system equilibrium is a function of the current extent of all active transporters, defined by

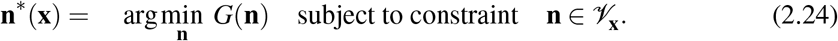

The space *V*_**x**_ can always be characterized by an equation of the form

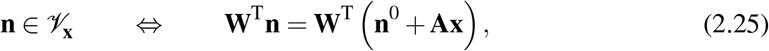

where the matrix **W** has *N* rows and *N* − dim(*V*_**x**_) columns, and the columns of **W** form a basis of 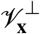.

#### A quasi-static simulation scenario

In this paper, the objective of the simulation is to evaluate the function **x** ⟼ **n**^∗^(**x**) when **x** describes a certain range. The expected result is an evolution of the quasi-static equilibrium state as active transporters work. In practice, when the input variable **x** is multidimensional, we make it follow a one-dimensional trajectory, so that the quasi-static equilibrium follows a one-dimensional trajectory in the space of states. Thus, simulations results look like an evolution as a function of a scalar variable. For instance, the example from Figure 1 involves two active transporters. If we run the simulation for **x**(*s*) = (*s, s/*2) with *s* describing the interval [0, *s*_max_], it means that the second pump works twice as slow as the first one. However, keep in mind that *s* does not represent time here, as our model does not include dynamics. By changing the way **x** depends on *s*, one can explore several possibilities of synchronization between active transporters.

Choosing a quasi-static scenario is a strong choice, as the individual dynamic of transporters is not taken into account, and transient regimes are not modeled. It corresponds to a limit case where the dynamics of passive transporters is very fast compared with the time span of the considered process, so that the system immediately goes back to an equilibrium when perturbed by active transport. Actually, assuming a quasi-static evolution boils down to separating transport processes into two categories: either the transport is slow, and it is not included in the model, or it is sufficiently fast, and it is considered instantaneous. However, our goal here is not to depict a time-dependent evolution. As the dynamic properties of transporters is not always well known, a quasi-static approach is a good way to highlight phenomena that do not depend on the transporter dynamics. Finally, the main advantage of choosing a quasi-static model is that it requires few parameters. While a dynamic simulation requires a function to determine the dynamics of each transporter, sometimes requiring to set many coefficients, the quasi-static model only requires the transporters’ stoichiometry, and the distinction between active and passive transporters.

### 2.3 Numerical implementation

In this technical section, we provide details about the numerical implementation used to run the simulations of the quasi-static scenario. Our simulation code^1^ is written in Python, and we leverage the automatic differentiation possibilities provided by the Jax library (Bradbury et al., 2018) to access the derivatives of *G* at least cost in terms of development. Using automatic differentiation is very convenient when working with variational problems, as resolution algorithms often require to evaluate the higher-order derivatives of energy or cost functions. Here, we were able to run simulations and perform sensitivity analysis with only a limited amount of calculations.

First, let us derive the nonlinear system that is actually solved in the numerical procedure. For a fixed input variable **x**, the corresponding equilibrium is defined by

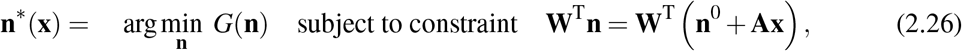

with the notation from (2.24) and (2.25). It can be shown that the solution to (2.26) satisfies some necessary first-order optimality conditions, also known as Karush-Kuhn-Tucker conditions (Nocedal and Wright, 2006, Section 12.3). These conditions read

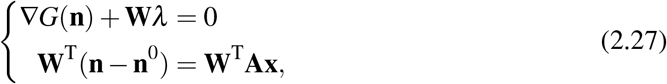

where *λ* is a Lagrange multiplier of size *N* − dim(*V*_**x**_). In (2.27), the first line describes the balance of forces that characterizes the equilibrium point. The multiplier *λ* is adjusted so that the artificial force **W***λ* enforces the constraint **n** ∈ *V*_**x**_. The second line corresponds to the constraint itself in matrix form.

To obtain a simpler system, we seek to eliminate *λ*. The matrix **W**, whose columns form a basis of 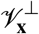, does not depend on **x**, as the *V*_**x**_ are parallel to each other. We define the matrix **V**, with *N* rows and dim(*V*_**x**_) columns, whose columns form a basis of *V*_**x**_. In particular, the columns of the block matrix (**V, W**) form a basis of ℝ^*N*^, and **V**^T^**W** = 0. By applying **V**^T^ to the first line of (2.27), we obtain the equivalent system of size *N*

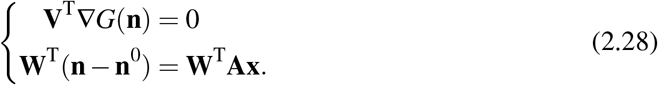

This system illustrates well that the coordinates of **n**^∗^(**x**) along directions collinear to *V*_**x**_ are defined by the balance of forces, while the coordinates along directions orthogonal to *V*_**x**_ are explicitly determined from **x**.

Let us write (2.28) in the case of the toy problem from Figure 3, with **V** and **W** defined by

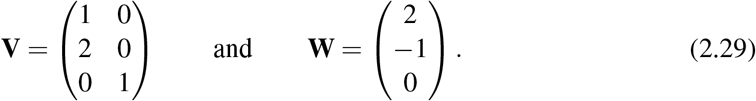

As the derivatives of *G*_elec_ and *G*_mech_ do not raise numerical issues, we only specify the expression for the chemical energy derivatives using (2.5), leading to the system

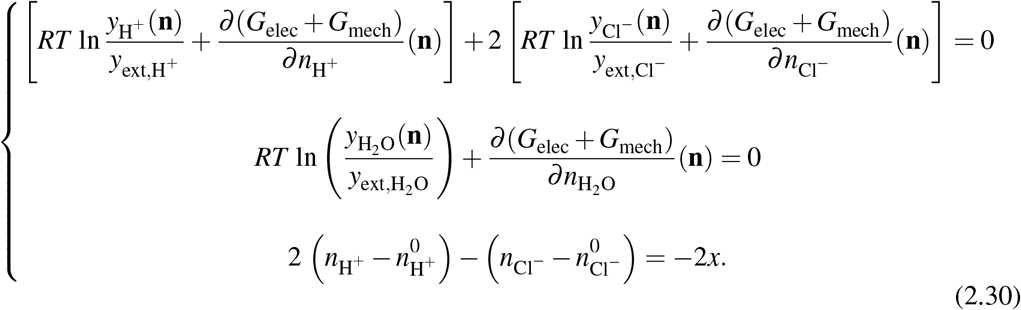

In practice, we aim to solve (2.30) using a Newton method (Nocedal and Wright, 2006, Algorithm 11.1). Here the presence of the logarithm function in the expression of chemical potentials represents a numerical difficulty, as the logarithm function is only defined for positive parameters, and takes large values for arguments close to zero. This may cause the Newton method to fail, either because the algorithm tries to evaluate the residual somewhere where it is not defined, or because the Jacobian matrix is poorly conditioned. To increase the numerical method robustness, we use a relaxed version of (2.30), following a so-called Cartesian representation technique, recently proposed by Jonval et al. (2025). Without entering too much into details, let us mention that, with this technique, the chemical potentials are represented as auxiliary variables 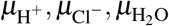 instead of functions of **n**. Then, *N* new lines are added to (2.30) to enforce a relation of the form *f* (*µ*_*i*,*A*_*/RT*) − *g*(*y*_*i*,*A*_(**n**)*/y*_ext,*A*_) = 0, where *f* and *g* are functions defined on ℝ and chosen to exhibit good numerical properties, namely

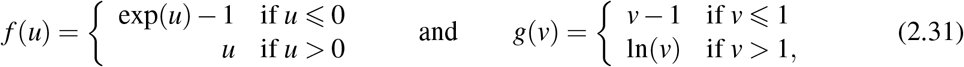

so that the enforced relation is either

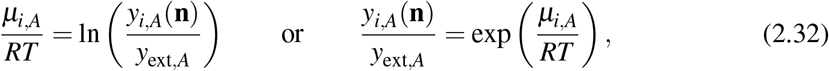

depending on the values of *y*_*i*,*A*_(**n**) and *µ*_*i*,*A*_ (note that *y*_*i*,*A*_(**n**) may take values out of [0, 1] if **n** has negative components). As a consequence, the relaxed system, which is actually solved in the numerical procedure, reads

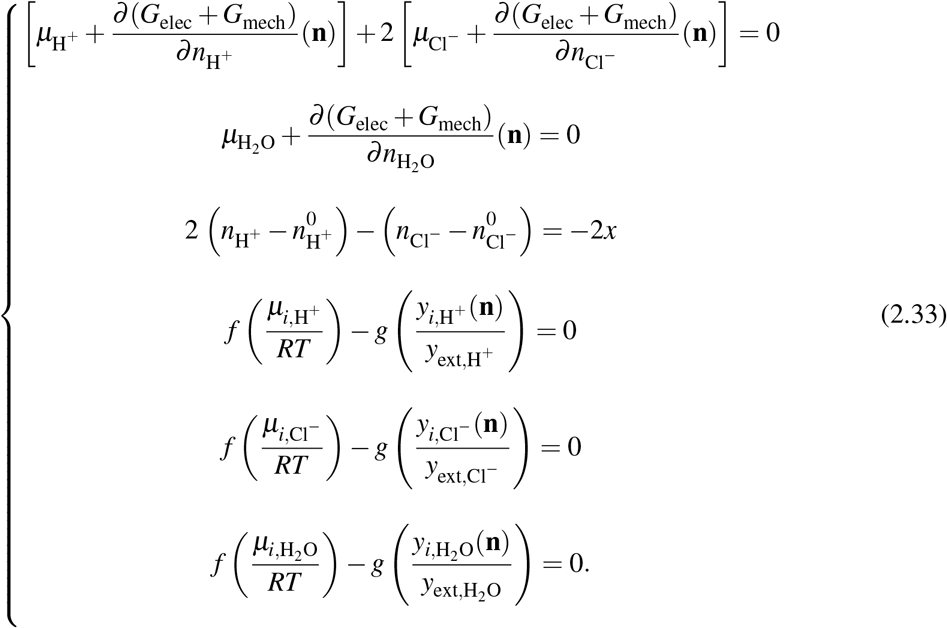

By solving (2.33) successively for *x* describing an interval [0, *x*_max_] we can compute the trajectory of the moving equilibrium with a limited risk of numerical instabilities.

In the next section, we apply the quasi-static formalism to perform a simulation involving the two-membrane complex from Figure 1 which symbolizes a guard cell.

## 3 Numerical simulations involving a simplified guard cell model

We now apply the energy-based formalism to a concrete system inspired from plant biology. The application case is the ion homeostasis in a guard cell during the stoma opening process. Located on plant leaves and stems, stomata are small pores that allow the plant to exchange water and carbon dioxide with the atmosphere. Each stoma is surrounded by two guard cells, that control its aperture by changing their shape. When the stoma opens, the change of the guard cell shape is only caused by its volume increase, due to water entering the guard cell. In this section, we simulate the cascade of transport processes, driven by two hydrogen pumps on the plasma and vacuole membranes, which leads to water being attracted into the cell. In the beginning of this section, we first describe the system we use to approximate the configuration of a guard cell, and we give a few details about the numerical procedure that we implemented. Then, we run a simulation and propose an interpretation of the results.

### 3.1 Description of a minimal model representing a guard cell

The processes leading to stoma opening involve the accumulation in the guard cell of different chemical species in a large amount, including ions and also sugars, to modify the osmotic pressure of the guard cell. The accumulation of ions and sugars depends on a wide variety of transporters localized in cellular membranes (Jezek and Blatt, 2017). Here, we adopt a minimal modeling strategy to build a model of ion transport in the guard cell, that features a limited number of transporters to remain interpretable. To be more specific, we limit the number of transporters and chemical species to those directly involved in the modification of the osmotic load. Therefore, we do not include calcium ions, since their function in guard cells is related to signaling and they do not contribute significantly to the osmotic effects (Li et al., 2024; Jezek and Blatt, 2017). During stoma opening, water uptake is mostly driven by the accumulation of potassium and anions like chloride and malate in the guard cells(Roelfsema and Hedrich, 2005). Chloride and malate play similar roles, electrical charge compensation and osmotic load, and therefore we only use chloride as a representative for the anions involved in this process. Last but not least, we include hydrogen and water in the model, as the former is transported by hydrogen pumps that drive the whole process, and the latter is directly responsible for cell volume changes. In summary, the simplified model involves four chemical species: hydrogen (H^+^), chloride (Cl^−^), potassium (K^+^), water (H_2_O).

The guard cell is represented by the two-membrane complex summarized in Figure 1. It contains two compartments, the cytoplasm (compartment 1) and the vacuole (compartment 2). In this context, the state vector is composed of the eight variables

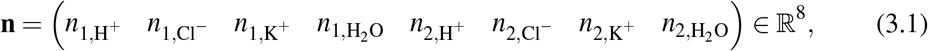

where the first four components are the chemical amounts in the cytoplasm and the other four are the chemical amounts in the vacuole.

The model includes a selection of the main transporters involved in the stoma opening process (see for instance Demes et al., 2020, Fig. 6), keeping in mind that chloride stands as a general anion here. The system includes one hydrogen pump on each membrane (i.e. the plasma and vacuolar membrane) with the plasma membrane pump ejecting hydrogen from the cytoplasm to the external environment (Merlot et al., 2007) and the vacuolar membrane pump ejecting hydrogen from the cytoplasm into the vacuole (Krebs et al., 2010). Further, each membrane includes a transporter coupling hydrogen and chloride. In the vacuolar membrane, the hydrogen/chloride transporter is an antiporter, as it transports the two species in opposite ways. The molecular identity of this antiporter is known (De Angeli et al., 2006; Jossier et al., 2010; Wege et al., 2014). In the plasma membrane, we introduce a chloride/hydrogen symporter (which transports both species in the same direction), that has not yet been molecularly identified (Guo et al., 2003; Jezek and Blatt, 2017). For this reason, its stoichiometry is not known. To obtain symmetric behaviors between transporters at the plasma and vacuolar membrane, we chose the following coefficients for this symporter: H^+^ + 2Cl^−^. Using different stoichiometric coefficients for the plasma membrane hydrogen/chloride symporter, such as H^+^ + Cl^−^ or H^+^ + 3Cl^−^, changes the amount of osmolytes imported in the cell per pump cycle, but it does not invalidate the analysis below. Finally, both membranes include a potassium channel, which is a passive transporter allowing this ion species to circulate, and water can freely circulate between compartments. For more details on ion transport in guard cells, see the review paper by Roelfsema and Hedrich (2005).

An advantage of using a quasi-static model is that it relies on few parameters, as all parameters associated with the transporter dynamics are not required. However, it is important to choose realistic values for the remaining parameters. The parameter values are gathered in Table 1. They mostly revolve around the geometrical properties of the cell model, as well as initial values for the chemical amounts and potentials. First, cell dimensions define the amount of water in each compartment. The guard cell is modeled as a cylinder that elongates in the longitudinal direction, where the diameter and initial length are about 10 *µ*m and 50 *µ*m, respectively (Meckel et al., 2007, Table 1). Following Mirasole et al. (2023, Figure 3), we assume that the vacuole accounts for 30 percent of the whole cell volume as the simulation begins. From these geometric parameters, we can evaluate the initial amount of water in both compartments.

**Table 1:** Numerical values of chosen parameters and initial conditions for the simulation.

| Parameter | Value |  |  |
| --- | --- | --- | --- |
| Cell diameter | 10 $\mu\text{m}$ | | |
| Cell wall surface stiffness | 30 Pa · m | | |
| Initial pressure | 0.5 MPa |  |  |
| Initial plasma membrane potential | -50 mV |  |  |
| Initial vacuole membrane potential | -40 mV |  |  |

| Parameter | External | Cytoplasm | Vacuole |
| --- | --- | --- | --- |
| pH | 5.7 | 7.1 | 5.8 |
| KCl concentration | 30 mmol/L | 15 mmol/L | 20 mmol/L |
| Compartment volume | — | 785 $\mu\text{m}^3$ | 3142 $\mu\text{m}^3$ |

Another parameter attached to the cell geometry is the stiffness of the wall, which is mainly a function of its Young modulus and its thickness. Several studies seem to agree around *E* ≈ 100 MPa for the Young modulus in the longitudinal direction and δ ≈ 0.1 − 1 *µ*m for the wall thickness (Woolfenden et al., 2017; Marom et al., 2017; Yi et al., 2018), which should result in a surface stiffness modulus *E*_s_ = *E ×* δ ≈ 10 − 100 Pa · m. In the computational model, we set the surface stiffness modulus to 30 Pa · m, which is consistent with previous studies while giving suitable volume changes in the simulation results.

We choose initial conditions in accordance with the literature. In their review, Jezek and Blatt (2017) gather values for ion concentrations and potentials for open or closed stomata. Following their numerical values, we set the initial pressure at 0.5 MPa (see also Franks et al., 2001), while the initial electric potential across the plasma membrane and the tonoplast are set to −50 mV and −40 mV, respectively, using the convention from Bertl et al. (1992). Concerning the initial concentrations for hydrogen, potassium and chloride, we set them to realistic values in the context of *in vitro* experiments involving guard cells (Demes et al., 2020; Mirasole et al., 2023).

### 3.2 Simulation of the guard cell during stoma opening

We now show simulations results involving the guard cell model from Figure 1. The system input is composed of the progress of each one of the two hydrogen pumps. These two input variables are stored in the vector **x** = (*x*_1_, *x*_2_) ∈ ℝ^2^, with *x*_1_ denoting the progress (in mol) of the plasma membrane pump, and *x*_2_ denoting the progress of the vacuole membrane pump. As mentioned in Section 2.2, *x*_1_ and *x*_2_ will both be functions of a scalar variable *s*, in order to obtain an evolution along a one-dimensional trajectory.

The first simulation involves a fixed ratio *α >* 0 between the two pump rates, i.e. **x**(*s*) = (*s, αs*). For each new value of *s*, the simulation computes a corresponding equilibrium state **n**^∗^(**x**(*s*)). Figure 5 shows the evolution of chemical amounts as a function of *s*, along with other physical quantities that can be computed from them. The figure includes pH values and the charge imbalance *Q*_*i*_(**n**) across membranes, while the hydrostatic pressure and electric potentials across membranes are defined by

**Figure 5:**
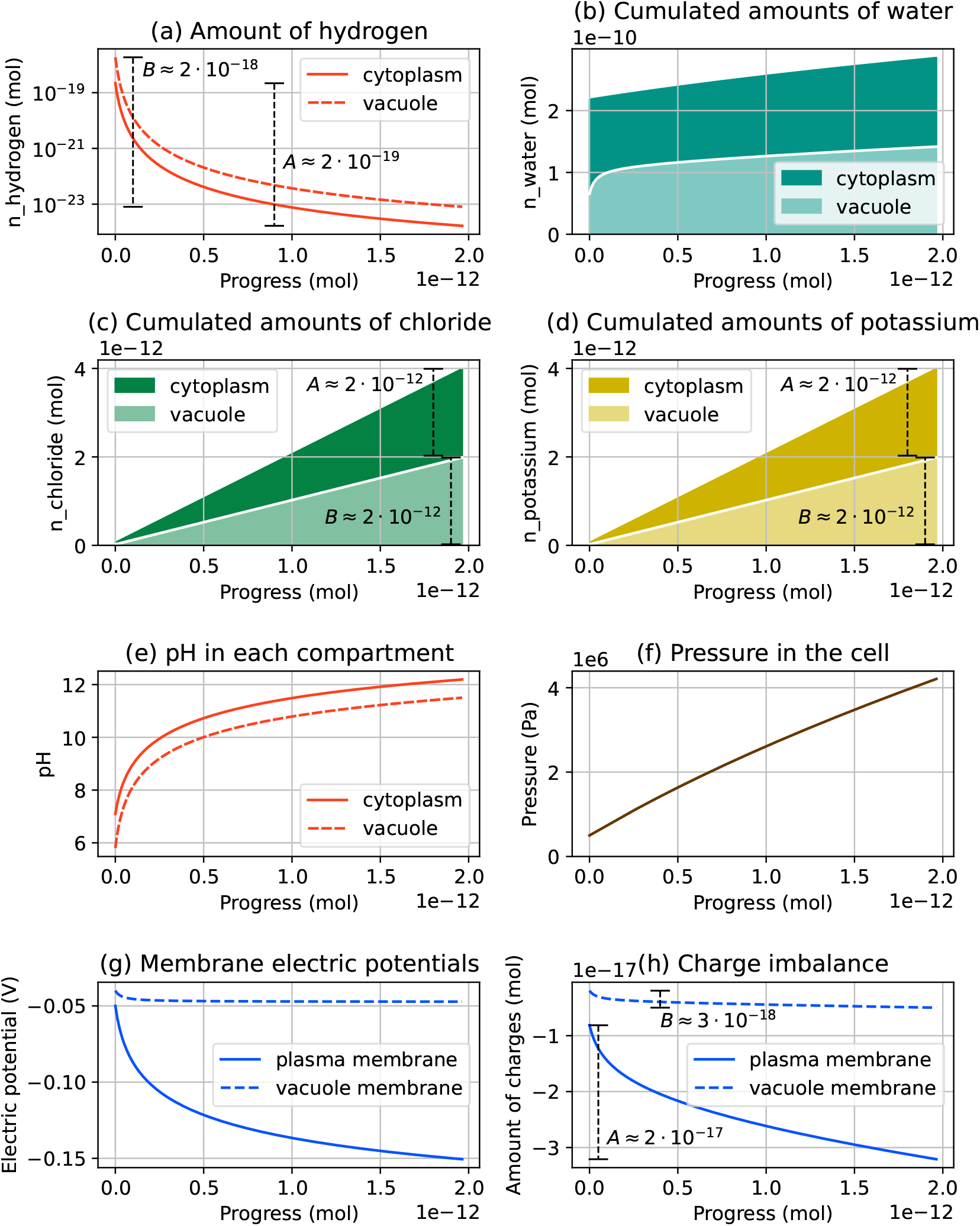
Simulation results for the guard cell model. The simulation output consists of the evolution of the amount of each chemical species as the progress of active transporters increases. Amounts of water, chloride and potassium are plotted in a cumulated way, with the upper part representing the amount in the cytoplasm and the lower part representing the amount in the vacuole. While pH reflects the hydrogen concentration, pressure and electric potentials are obtained from derivatives of the energy function.

**Figure 6:**
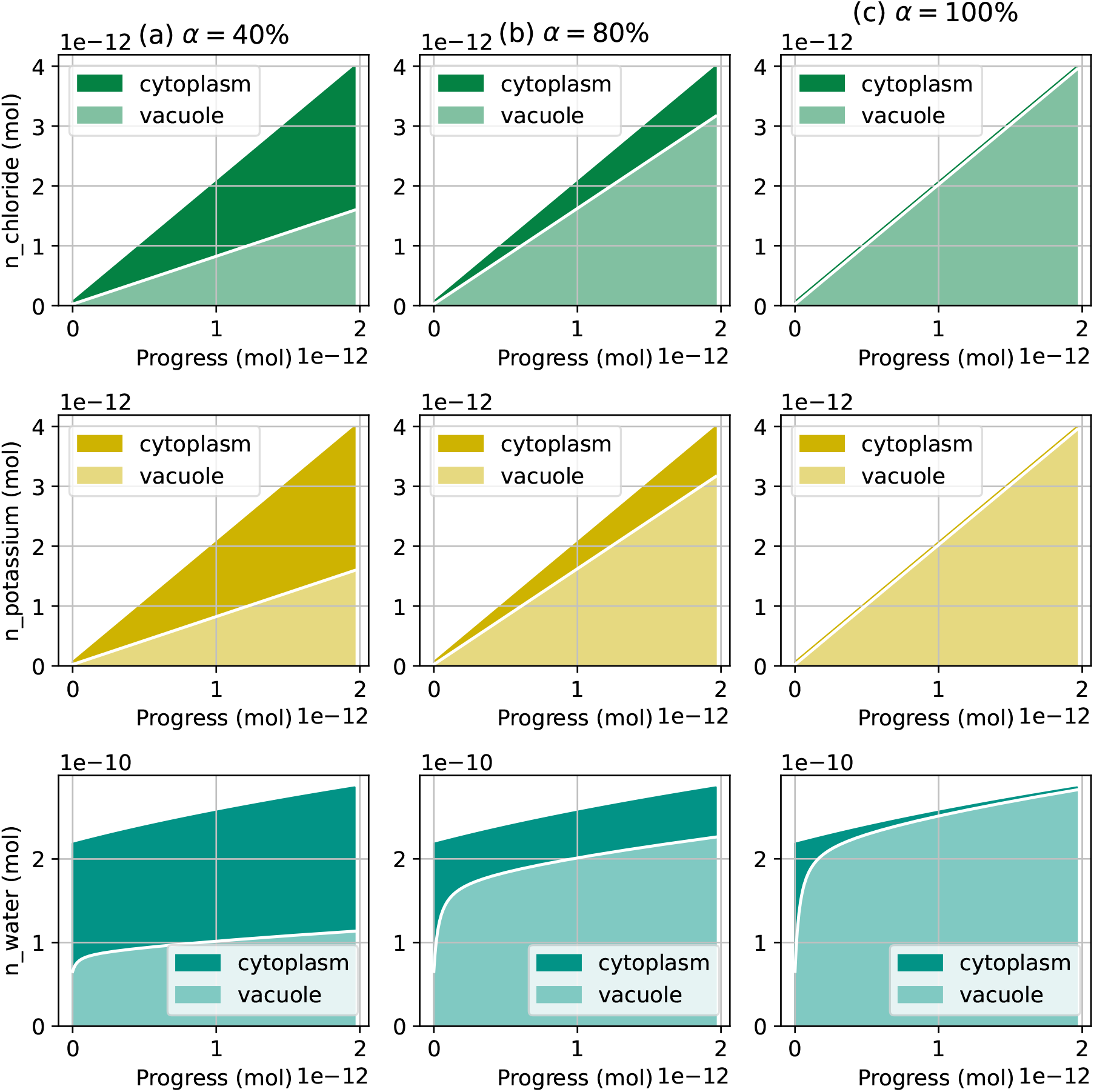
Simulation results with constant ratios between pump rates. The rates of the vacuole pump are *α* = 40% (left), *α* = 80% (middle), and *α* = 100% (right). The plotted values are the amounts of chloride (top), potassium (middle) and water (bottom). All plots on a same row share the same *y*-axis. The ratio between osmolyte and water transport at the vacuole membrane and at the plasma membrane is asymptotically similar to the ratio between the hydrogen pump rates.

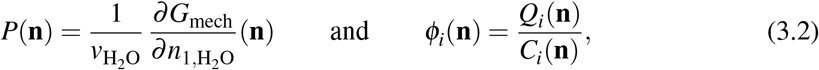

respectively (see (2.13) and (2.9) for notations). Electric potentials are plotted following the sign convention from Bertl et al. (1992), i.e a negative potential value across a membrane means that the cytoplasm is at a lower electric potential than the other compartment. The simulated system generally follows the biologically expected behavior, as the activation of the pumps results in chloride, potassium, and finally water entering the cell. Also, we notice that the range of variation of electrical potentials and turgor pressure are in the right order of magnitude (Franks, 2003; Woolfenden et al., 2017). The behavior of hydrogen concentrations, in contrast, is less expected. While the pH usually remains stable in the cytoplasm and slightly acidic in the vacuole, in our simulations both compartments become strongly basic as the cell gets emptied of its hydrogen. This point is addressed in the next section. However, it should be noted that the hydrogen depletion is very slow. At the end of the simulation, the amount of hydrogen in the cell has decreased by 2 · 10^−18^mol, while the amount of hydrogen actively ejected by the pump is 2 · 10^−12^mol.

In this configuration, numerical experiments suggest that the plasma membrane pump drives ion and water transport across the plasma membrane, while the vacuole membrane pump drives exchanges between the cytoplasm and the vacuole. For instance, in the results from Figure 5, the ratio between pump rates is *α* = 50%, which means that when the plasma membrane pump ejects 100 protons from the cell, the vacuolar pump sends 50 protons into the vacuole. As a result, 50% of the osmolytes entering the cell are sent to the vacuole, which ends up representing 50% of the whole cell volume. To confirm this trend, we run the same simulation with other values for *α*, namely 40%, 80% and 100%. We do not explore values of *α* greater than 100%, as they raise numerical instabilities due to chemical amounts in the cytoplasm becoming close to zero. Results, gathered in Figure 6, show that, at the end of the simulation, the fraction of solute and water in the vacuole asymptotically matches the ratio between the pump rates. If the stoichiometric coefficients of the hydrogen/chloride symporter in the plasma membrane were changed, then the final amount of osmolytes in the vacuole would not be the same, but it would still be proportional to the vacuolar membrane pump rate. Note that pH, electric potential and pressure values do not change significantly between cases from Figures 5 to 7.

Therefore, the fraction of the cell volume occupied by the vacuole can be controlled through the ratio between the rates the two pumps. When inducing stoma opening in *in vitro* guard cells, Mirasole et al. (2023) observed that the cytoplasm volume remains approximately constant along the process. To reproduce this phenomenon in the simulation, we set as previously the plasma membrane pump rate at 1 (i.e. *x*_1_(*s*) = *s*), and then we adjust the vacuole pump progress *x*_2_(*s*) at each simulation step, so that the amount of water in the cytoplasm remains constant. The results of the simulation with constant cytoplasm volume are shown in Figure 7. The vacuole pump rate 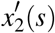 can be read as the slope of the vacuole pump progress in Figure 7b. For each value of the progress *s*, it is equal to the current fraction of cell volume occupied by the vacuole (Figure 7a). Initially at 30%, this fraction has increased to 50% by the end of the simulation, while the slope of the vacuole pump progress graph is slightly larger at the end of the simulation than at the beginning.

**Figure 7:**
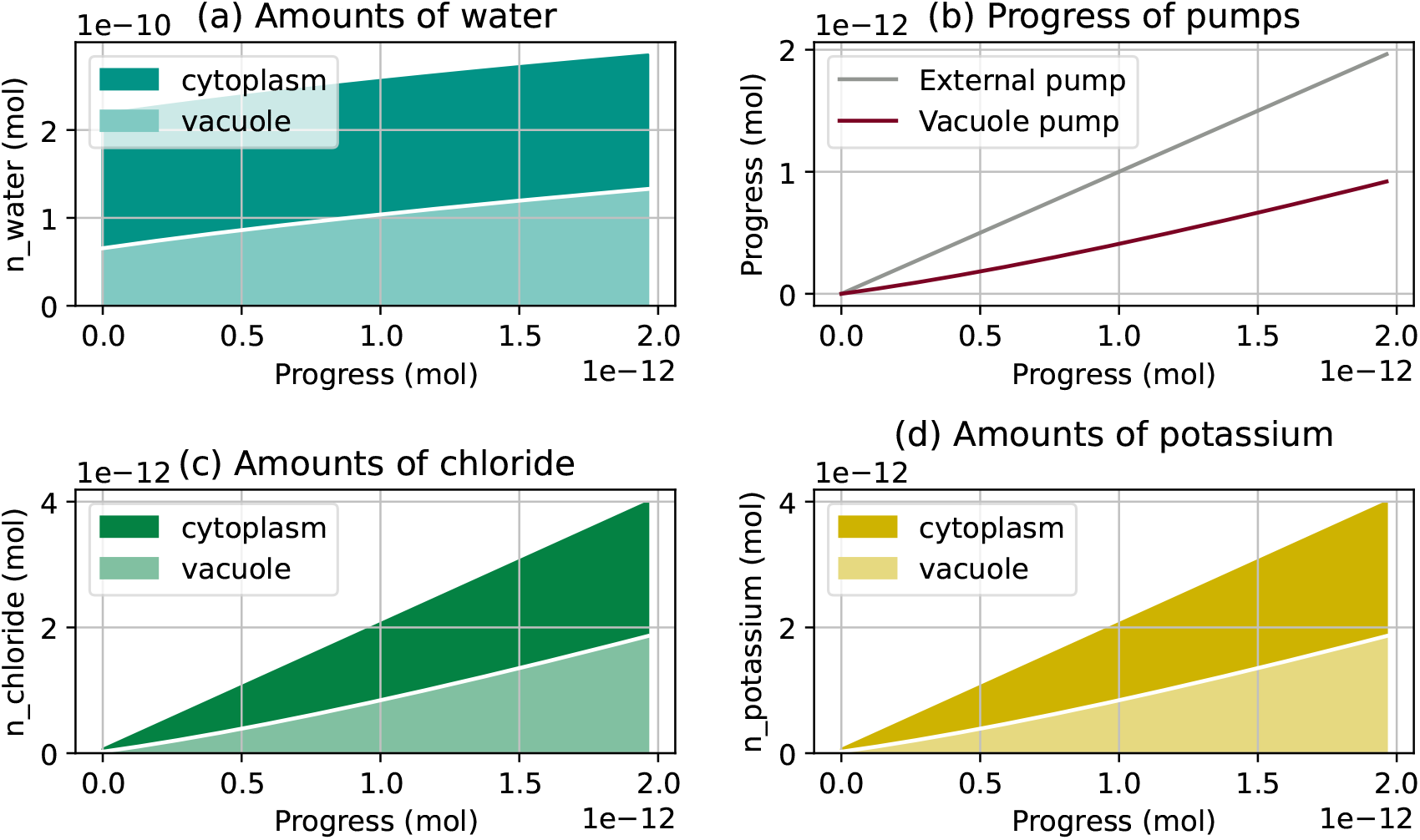
Simulation result involving a constant-volume constraint. The rate of the vacuole membrane pump is controlled to keep the cytoplasm volume constant. The plot (b) shows the progress of both pumps along the simulation.

Because the transport that occurs at one membrane does not influence the transport at the other membrane, it is possible to describe separately the events happening at each membrane, using Figure 1. At the plasma membrane, hydrogen expelled from the cell by the pump is almost completely compensated for by hydrogen entering back through the symporter, accompanied by chloride. This explains why the amount of chloride in the cell is approximately twice as large as the pump progress *s* (Figure 5c). However, this cycling is not perfect, and the amount of hydrogen entering the cell through the symporter is slightly smaller than that ejected by the pump. Consequently, there is a net loss of hydrogen by the cell. (Figure 5a). The massive intake of chloride is accompanied by an approximately equal intake of potassium (Figure 5d), through the potassium channel. As these two ions represent most of the charge transfer, the difference between chloride and potassium intake can be read in the charge imbalance plot (Figure 5h). Finally, water intake (Figure 5b) is synonymous with volume increase, and the reaction of the elastic cell wall to this deformation can be seen in Figure 5f through the pressure increase. A similar chain of events occurs at the vacuolar membrane, except that hydrogen is pumped into the vacuole, and it goes back into the cytoplasm through the antiporter, in exchange for chloride entering the vacuole. Again, the cycling of protons is not ideal and there is a net loss of hydrogen from the vacuole. This, together with the loss of hydrogen at the plasma membrane, results in an increase of the cell pH. An interpretation of the physical processes causing this evolution is proposed in Section 4.3.

### 3.3 Impact of hydrogen buffering mechanisms on simulation results

So far, we have considered a model involving solute and water transport only, without chemical reaction. While simulation results show a plausible evolution for most physical quantities, a discrepancy remains between the simulation and experimentally observed guard cell behavior. In the simulation, hydrogen tends to disappear from the cell, leading to a very basic pH in both compartments. On the other hand, experimental observations suggest that the cytoplasm pH remains close to 7, while the vacuole pH remains acidic or slightly increases, depending on reports (Andrés et al., 2014; Li et al., 2024; Cha et al., 2024). A key parameter in the cell, pH is actually well controlled using a range of mechanisms, called hydrogen buffering. These mechanisms are able to capture, release or transport hydrogen, to maintain the pH in a desired range in each compartment (Kurkdjian and Guern, 1989). To remedy this inconsistency in our simulation, we now include a hydrogen buffering mechanism in the model, and we investigate its consequences on the simulation output. Though both the cytoplasm and the vacuole of guard cells feature hydrogen buffering in a guard cell, the vacuole pH is less tightly controlled compared with the cytoplasm pH (Andrés et al., 2014; Li et al., 2024; Cha et al., 2024). For the sake of simplicity, we only consider hydrogen buffering in the cytoplasm.

In reality, hydrogen buffering results from several complex mechanisms, including chemical components (e.g. organic acids, phosphates), metabolic activity, solute transport and buffering capacity of the cell wall (Kurkdjian and Guern, 1989; Poznanski et al., 2013). In the model, we summarize these mechanisms in the form of a chemical reaction that compensates the variations of hydrogen concentration in a compartment. We represent buffering as an acid-base reaction of type

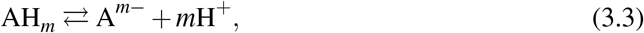

where A is a species able to bind with hydrogen. When the hydrogen concentration decreases, the reaction provides some hydrogen by consuming AH_*m*_, while the reaction produces AH_*m*_ when hydrogen is present in excess. The stoichiometric coefficient *m* is introduced as a numerical tool to adjust the strength of pH regulation. We choose a large *m* to create a strong buffering capacity in the cytoplasm, without inducing perturbations of osmotic effects due to the presence of large amounts of buffer molecules in the compartment. Here, we choose *m* = 10^4^ and we set the initial concentration of the buffering solution to 1 mmol*/*L in the cytoplasm. In the same fashion as passive transporters, this chemical reaction is represented by one of the vector directions that span *V*_**x**_, as it gives the system the freedom to evolve toward production or consumption of AH_*m*_.

Figure 8 shows the simulation results for the buffered model, with the same pump rates as in Figure 5, i.e. 1 for the external pump and 0.5 for the vacuole pump. The hydrogen amounts evolve differently compared with previous simulations. Here, the cytoplasm pH remains stable and close to 7, and the vacuole pH remains in acidic values, which is in agreement with experimental observations. Note that the pH in the vacuole decreases more than what is suggested by experimental measures. The amplitude of pH variations in the vacuole can be corrected by adding a buffering mechanism in this compartment. Interestingly, the evolution of chloride and potassium amounts is also changed, and the connection between compartment volumes and pump rates observed in Figure 6 is now more loose. While, without buffer, each pump is responsible for making osmolytes cross the membrane it is located on, the introduction of the buffering mechanism makes the effect of both pumps more intricate. Our simulations suggest that the vacuolar membrane pump triggers osmolyte transport all the way from the external environment into the vacuole, and the amount of osmolytes crossing each membrane is not proportional to the pump rates anymore. Though studying the buffered system in details is out of the scope of this article, results from Figure 8 show that our model can be extended to produce more realistic simulations.

**Figure 8:**
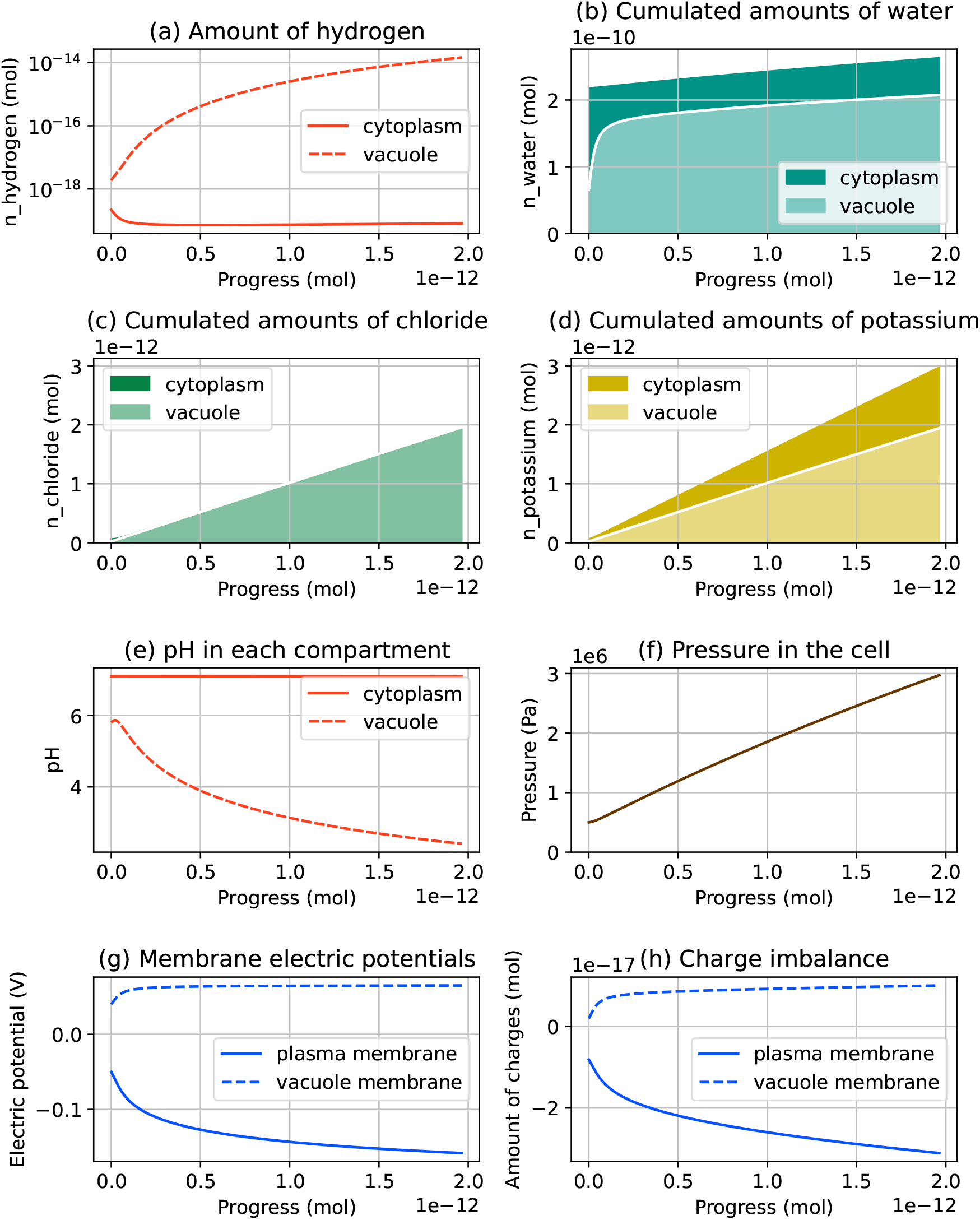
Simulation result for the guard cell model with hydrogen buffer. The plotted physical quantities are the same as in Figure 5. In particular, the cytoplasm pH values follow a more realistic behavior when hydrogen buffering is taken into account.

## 4 Sensitivity analysis using second derivatives of energy

In this section, we perform a sensitivity analysis of the guard cell model, based on second-order information about the energy function. More specifically, our aim here is to explain the range of variation predicted by the model for all variables, and also to highlight the couplings between variables created by the different terms of the energy function. After a short calculation that emphasizes the role of the energy second derivatives in determining the system response to a perturbation, we give some insight about the physical meaning of the energy second derivatives.

Finally, we implement this reasoning by comparing second derivative coefficients and simulation results in the case of the guard cell model. As *G* is a regular function of several variables (*n*_1_, *n*_2_, …), its second derivatives at **n** are stored in a symmetric matrix called the Hessian matrix, denoted by ∇^2^*G*(**n**). In particular, the Hessian coefficient at index (*i, j*) is denoted by 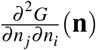.

### 4.1 Second derivatives of the energy function: a physical analogy

First, let us clearly identify where the energy second derivatives appear in the calculations. Remember that, for a fixed **x**, the equilibrium state **n**^∗^(**x**) is solution to the nonlinear system (2.28). Now, assume that **x** is subject to a perturbation d**x**. The equilibrium perturbation d**n**^∗^ is the solution to a tangent linear system involving the Hessian matrix of the energy function. This linear system, obtained by differentiating (2.28), reads

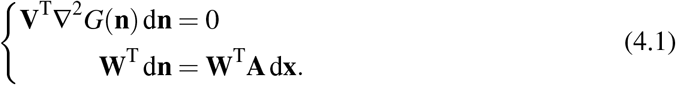

Like previously, the first line of (4.1) concerns the coordinates of d**n** in the direction of *V*_**x**_, which adjust to re-establish a physical equilibrium, while the remaining coordinates are explicitly determined from d**x** by the second equation. The Hessian matrix plays a role in the evolution of **n** in the directions of passive transporters, as a reaction to an imposed change in the directions orthogonal to *V*_**x**_. While forces are defined by the first derivative of the energy, this analysis explains why second derivatives must be considered in a sensitivity analysis as they reflect force responses to perturbations. This idea can be illustrated through a simple physical analogy. Consider a spring of stiffness *k*, clamped at one extremity and subject to a force *F* at the other extremity, causing it to extend by a length *x* in a one-dimensional way. The spring elastic energy function reads

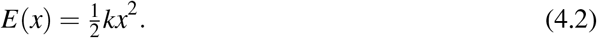

Also, *E*^*′*^ and *E*^*′′*^ denote the first and second derivatives of the single-variable function *E*. Here the second derivative of *E* is the stiffness *k*, which defines the spring resistance to changes of lengths. If the force applied by the external user at the spring extremity varies from *F* to *F* + Δ*F*, the spring length changes by Δ*x* = Δ*F/k*. In other words, the spring stiffness gives an indication about how much the system resists to perturbations. In the same fashion, if we consider a nonlinear spring with an energy function *E*, the force applied at its extremities satisfies *F* = *E*^*′*^(*x*), and as a consequence, a force perturbation Δ*F* causes the length to change at first order by Δ*x* = Δ*F/E*^*′′*^(*x*), where *E*^*′′*^(*x*) plays the role of an immediate stiffness.

The second derivatives of *G* are like the stiffness of a spring. However, as opposed to the spring energy function, *G* is a function of several variables, and its second derivatives are gathered in the Hessian matrix ∇^2^*G*(**n**). Diagonal terms in this matrix are stiffness-like coefficients. For a given variable *n*_*i*,*A*_, they define to which extent a variation of *n*_*i*,*A*_ results in a force causing *n*_*i*,*A*_ to go back to equilibrium. On the other hand, nondiagonal terms denote couplings between the different variables. The coefficient *∂* ^2^*G/*(*∂n*_*i*,*A*_*∂n*_*j*,*B*_)(**n**) denotes to which extent a variation of *n*_*i*,*A*_ results in a force causing the variation of *n* _*j*,*B*_. An example of such coupling appears when a transfer of charged particles affects charged particles from another species.

Pushing the spring analogy a little further, we now consider *n* springs in series. The springs chain is clamped at both end, so that its total length is imposed. At equilibrium, the individual deflection of the *i*-th spring is denoted by *x*_*i*_, and due to the imposed total length, the spring deflections all add up to an imposed value *y*. The *x*_*i*_ values are determined by solving the minimization problem

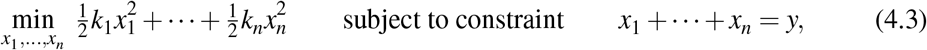

where *k*_*i*_ is the stiffness of the *i*-th spring. Now, if the imposed cumulated deflection *y* changes by a perturbation Δ*y*, the system will find a new equilibrium state, where each deflection *x*_*i*_ has changed by a small displacement

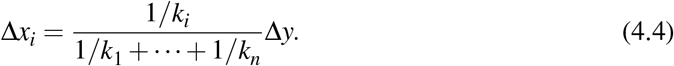

The sensitivity of each spring to the perturbation is inversely proportional to its stiffness. If the first spring has the largest stiffness, then its deviation from its initial state is smaller than the deviation of other springs, but it is also even smaller as there are many springs in the chain to absorb the perturbation. In other words, we could say that the initial state of springs with a larger stiffness is preserved *at the expense* of that for springs with a lower stiffness. Similarly, if each spring is nonlinear and characterized by its energy function *E*_*i*_(*x*_*i*_), then, solving a minimization problem similar to (4.3) yields the first-order responses to a perturbation Δ*y*,

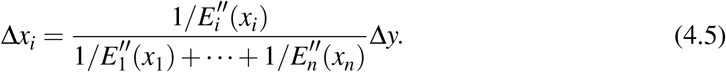

The balances that exist in our system, such as the balances of concentrations, of charges, or the mechanical equilibrium of the system, can be compared with the springs in series. Each part of the energy function promotes one balance, taking higher values when the balance is not respected. But, somehow, all these balances are in competition with each other, and, when a perturbation occurs, the term with the largest second-order derivative wins, i.e. the corresponding balance is more preserved, at the expense of other balances. In the remaining of this section, we exploit this analogy to interpret the simulation results from Section 3.

### 4.2 Comparison between Hessian coefficients and simulation results

In this section, we compare the energy Hessian coefficients with the range of variation of chemical amounts in the simulation results. While this analysis is straightforward for systems involving a few variables, the exercise becomes increasingly difficult as the model gets complex. When more variables are added, Hessian matrices become larger. To illustrate the role of Hessian coefficients, we focus on the guard cell model without hydrogen buffering, as it features fewer variables and is easier to interpret the model with hydrogen buffering.

Figure 9 shows the Hessian matrix for each term of the energy function, along with the Hessian matrix for the total energy function. All these Hessian matrices are evaluated at the initial state **n**^0^, but the trends highlighted by our sensitivity analysis appear to remain valid for the whole simulation (see Hessian matrices evaluated at the final state in S1 Fig.). Note that the values written in Figure 9 are in logarithmic scale, i.e. 22.6 on the matrix heatmap means that the corresponding coefficient is equal to 10^22.6^ J*/*mol^2^ in absolute value, while a blank box denotes a null coefficient. Therefore, the numbers visible in the total Hessian heatmap are not the sum of the corresponding numbers in the other heatmaps. Often, a coefficient in the total Hessian matrix (Figure 9d) approximately exhibits the same order of magnitude as the largest corresponding coefficient in the three other matrices. Concerning the axis labels of the figure, each row and column of a Hessian matrix is associated with one coordinate of **n**. Chemical amounts are organized in the same order as in (3.1), i.e. amounts in the cytoplasm first and amounts in the vacuole next. For instance, the coefficient at row 2 and column 7 corresponds to the coupling between 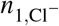 and 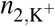.

**Figure 9:**
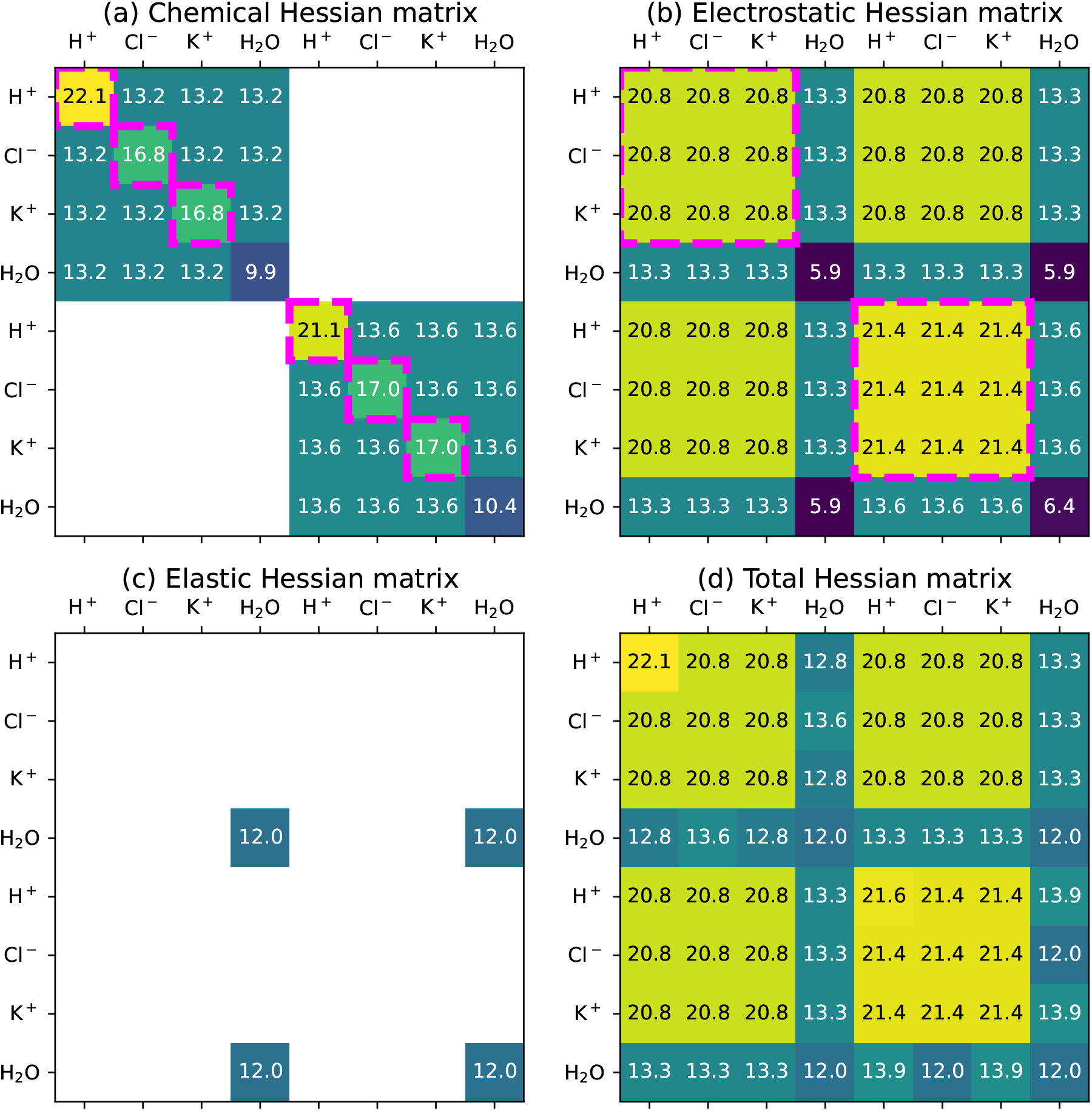
Hessian matrices for each term of the energy function, evaluated at the initial state of the simulation. Hessian coefficients (in J*/*mol^2^) are represented in logarithmic scale, i.e. 22.6 means 10^22.6^ and a blank case means that the coefficient is zero.

We now extract information from the Hessian matrices, and compare the magnitude of their coefficients with the variations of variables in Figure 5. As explained above, while Hessian coefficients play the role of spring stiffnesses, the amplitude of variations for associated variables denotes to which extent they deflect from their initial value.

First, the chemical energy Hessian matrix (Figure 9a) is made of two diagonal blocks, as the chemical energy function is composed of two terms (*i* = 1, 2 in (2.3)), with each term depending only on chemical amounts in one compartment. In each block, the diagonal coefficients associated with the amounts of hydrogen, chloride and potassium (see framed coefficients in the heatmap) are much larger in magnitude than nondiagonal parameters. The coefficient associated with hydrogen is approximately 10^9^ times larger than nondiagonal parameters, while the coefficients associated with chloride and potassium are ∼ 10^3^ times larger. From this remark, we can understand that the chemical energy term mainly acts on the individual evolution of each one of these amounts of substance, without creating a significant coupling between ion species. Moreover, in each compartment, the diagonal coefficient associated with hydrogen is much larger than that associated with chloride and potassium. It can actually be derived from (2.6) that these diagonal coefficients are inversely proportional to the amounts of the corresponding species (from the derivatives of the log function). In other words, as hydrogen is present in much smaller amount than other ions, a variation of its amount has a larger impact on the chemical equilibrium than the same variation for a more abundant species.

This difference between coefficients has its counterpart in Figure 5 (a, c, d), where the annotations indicate the variations range of each chemical amount (in mol) in the cytoplasm (*A*) and in the vacuole (*B*). In particular, the variations of hydrogen amounts (*A* ≈ 10^−19^ mol, *B* ≈ 10^−18^ mol) are several orders of magnitude smaller that variations of chloride and potassium (*A* ≈ *B* ≈ 10^−12^ mol), which can be explained by the hydrogen variations being highly restricted by a stronger feedback force. Comparing the variations of the same physical quantity between the two compartments is less straightforward. The variation range of hydrogen amounts in both compartments is mostly a consequence of initial conditions, without significant impact of the pump rates. On the other hand, we saw in Figure 6 that the ratio between chloride and potassium quantities in both compartments can be fully explained by the ratio between the pump rates.

The chemical Hessian matrix does not exhibit strong diagonal coefficients for water. From the point of view of water, the main impact of chemical forces is the coupling with ion species, represented by nondiagonal coefficients (10^13.2^). Adding some ions into a compartment creates a force attracting water, which is the osmosis effect. As expected, this coupling does not depend on the ion species. The term that offers the most resistance to variations of the water amount is the elastic energy term, as it exhibits the largest stiffness-like coefficient for water (10^12^, see Figure 5c). As a consequence, the volume of water entering the cell as a result of osmotic pressure can be entirely adjusted in our simulation, just by tuning the cell wall stiffness parameter, with a negligible impact on other variables.

The electrostatic Hessian matrix (Figure 5b) is more difficult to read, due to coupling between compartments. The electrostatic energy expression (2.9) falls into two terms. One term, associated with the plasma membrane, depends on amounts in both compartments, while the other term, associated with the vacuole membrane, only depends on amounts in the vacuole. In the matrix, the three identical 4 *×* 4 blocks are due to the first term, while the bottom right block, different from the others, includes in addition second derivative information from the second term. Interestingly, the electrostatic energy term creates couplings between water and charged species, as shown by the nonzero coefficients in columns and rows corresponding to water. Though water is not charged, the variations of water amounts cause variations in the cell geometry and in the electric capacitance of membranes. Removing water from a compartment tends to decrease its membrane area, making the capacitance smaller and the electric potential larger. But the most significant coupling generated by electrostatic effect concerns, as expected, charged species. This coupling corresponds to the framed zones in the electrostatic Hessian matrix. Even though it does not appear clearly on the heatmap, we know from (2.9) that the electrostatic energy is connected to the amount of charges in each compartment, denoted by 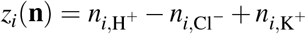. It takes large values when charge imbalance occurs at one membrane, i.e. a difference of potential builds across the membrane, but it is indifferent to everything that leaves *z*_1_(**n**) and *z*_2_(**n**) unchanged. The coefficients visible in the electrostatic Hessian matrix denote the sensitivity of the forces that promote the balance of charges on each side of both membranes.

To measure the discrepancy between the system state and a balance of charges, we plotted in Figure 5h the charge imbalance across each membrane, defined by *Q*_*i*_(**n**)*/F* (see 2.9). It denotes the amount of charges (in mol) stored in each membrane/capacitor (see the circuit analogy in Figure 2). Like previously, the annotations show the charge imbalance variations at the plasma membrane (*A* ≈ 10^−17^ mol) and at the vacuole membrane (*B* ≈ 10^−18^ mol). Because charge imbalance shares units with chemical amounts, its variations (Figure 5h) can be compared with the amount variations of all reactants (Figure 5a, c, d). Variations of the charge imbalance at the plasma membrane are slightly larger in amplitude than variations of the amount of hydrogen in the cell, and much smaller than variations of the amount of chloride and potassium in the cell (10^−18^ mol *<* 10^−17^ mol ≪ 10^−12^ mol). This is consistent with Hessian matrices, as the coefficient for electrostatic coupling in the boxed zone (10^20.8^) is slightly smaller than diagonal coefficients associated with hydrogen (10^22.1^, 10^21.1^) in the chemical Hessian matrix, and much larger than diagonal coefficients associated with chloride and potassium (10^21.1^ *>* 10^20.8^ ≫ 10^17^). Concerning charge imbalance at the vacuolar membrane (*B*), simulations show that it is strongly impacted by a combination of initial conditions and pump rates, and it cannot be explained using Hessian coefficients only.

The connection between the Hessian matrix coefficients and the amplitude of variation of the system variables illustrates well the role played by the second derivatives of the energy function. Like springs with a large stiffness, balances associated with large Hessian coefficients (hydrogen, charges) tend to resist to a perturbation, while balances associated with small Hessian coefficients (chloride, potassium, water) absorb most of it. However, even springs with a very large stiffness deflect a little when subject to a perturbation, and so does the amount of hydrogen, resulting in the hydrogen loss observed in Figure 5a.

Though the system evolution is also influenced by other parameters such as pump rates and transport stoichiometry, studying the Hessian matrix of each energy term allows us to identify trends in the range of variation of physical quantities. Note that these trends only depend on the energy function and the current system state, and they do not depend on the nature of transporters. As a consequence, this analysis remains valid for similar systems involving embedded compartments and osmolyte transport through membranes, including systems where the disposition of transporters is not known with certainty.

### 4.3 An interpretation for turgor pressure building in plant cells

Comparing the sensitivities of all the balances that govern the system allows to qualitatively understand the hierarchy between the physical phenomena at play, and explain why a property is enforced at the expense of another one. In particular, we can interpret the guard cell scenario from Section 3 through the prism of this hierarchy between the terms of the energy function, going from the activation of hydrogen pumps all the way to water entering the cell. Our simulation results in Section 3.2 suggest that transport processes on both membranes function independently of each other, each pump driving all transport across its membrane. For this reason, we show in Figure 10 an interpretation for solute transport at the plasma membrane, while a similar summary can be made for the vacuole membrane transport. In this cascade of transport reactions, each step restores one balance by perturbing another one that is lower in the hierarchy. For instance, in Figure 10, expelling hydrogen from the cell results in hydrogen entering back through the symporter, accompanied by chloride, as deflecting from the chloride chemical balance is much cheaper than deflecting from the hydrogen chemical balance. In the same fashion, the amount of potassium is forced to stick to that of chloride to restore the balance of charges, as this balance is much more sensitive than that of potassium. Membrane polarization, characterized by an electric potential building across a membrane, reflects the deflection of the system from electric neutrality. Finally, the resulting imbalance in chloride and potassium is compensated for by water entering the cell. The final volume change is a compromise between chemical forces and mechanical forces, and it mostly depends on the mechanical parameters of the cell wall. Note that, if potassium could not enter the cell to ensure an electrically neutral flow of osmolytes, a small amount of chloride would still enter the cell at the expense of the charge balance, as the balance of hydrogen is more sensitive than the balance of charges.

**Figure 10:**
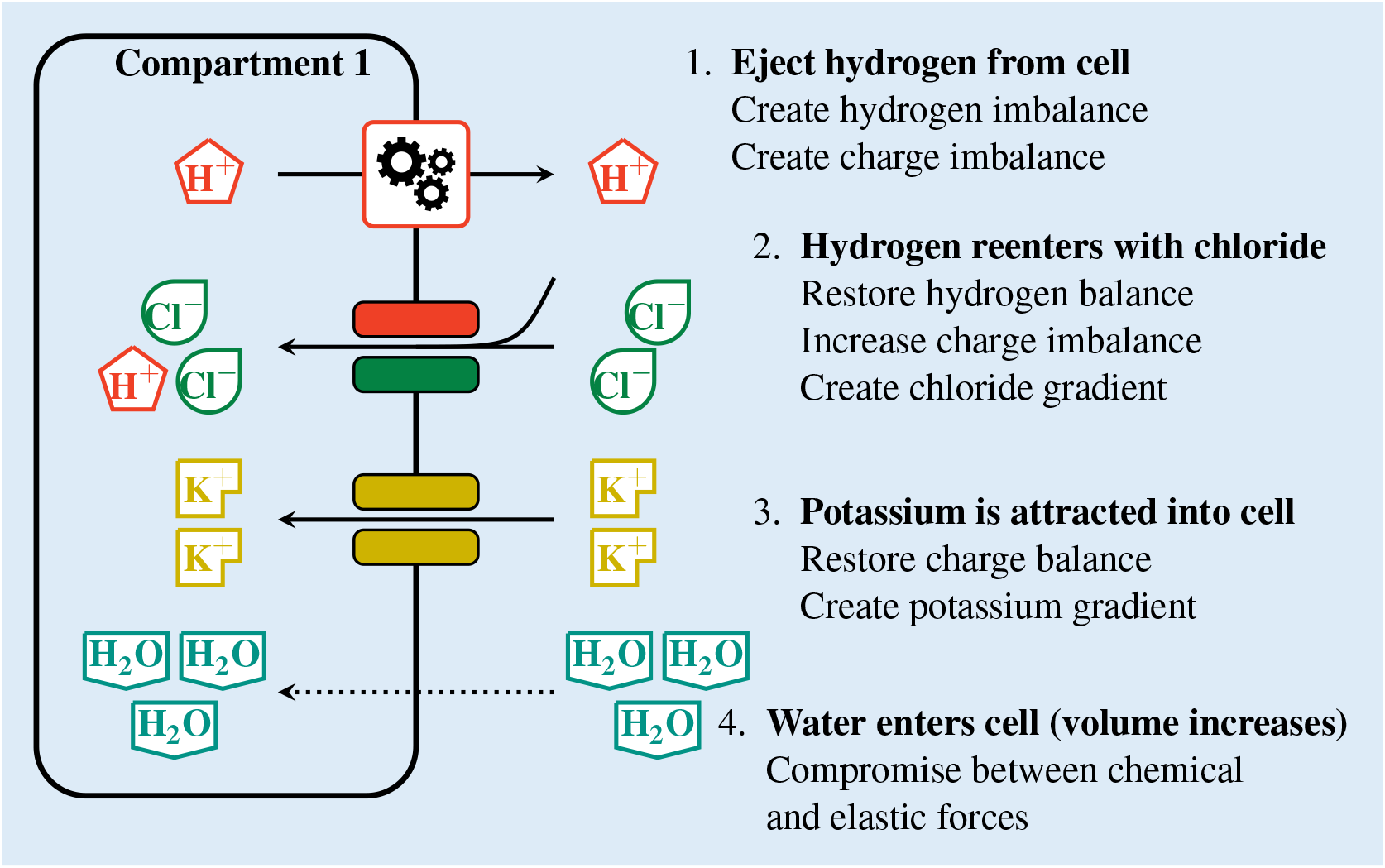
Summary of the transport cascade at the plasma membrane in the guard cell model without buffering. The vacuole is not represented in the figure. In steps 2 and 3, one balance is restored at the expense of another less strictly enforced balance.

The difference of sensitivities illustrates well how, by acting on hydrogen, which is present in very small amount, the cell manages to transfer into its vacuole an amount of water that is several orders of magnitude larger than that of hydrogen. Generating a perturbation on a strictly enforced balance creates a leverage effect. As it is diverted toward other balances, the perturbation gets amplified and involves larger-scale transport, which remains cheaper than maintaining a hydrogen imbalance. Even though transporters play a key role in determining the possible directions of evolution for the system state, the hierarchy between all forces, as well as involved orders of magnitude, are mostly a matter of energy function.

## 5 Discussion and Conclusion

### Summary

In an attempt to simplify the modeling of multi-physics couplings in multi-compartment plant cell physiology, we proposed in this paper a model based on an energy function integrating electrochemical and hydro-mechanical effects. We applied this energetic approach to study solute and water exchanges in a guard cell during stoma opening. By using a quasi-static scenario, we investigated the system properties without relying on transporter dynamics, thus reducing the number of parameters to set and the model complexity. We performed some simulations on the guard cell model with and without hydrogen buffering mechanisms, along with an analysis of the system response to perturbations. We showed that this model can perform biologically plausible simulations, as well as provide some insight about the role and influence of each physical force in the system.

### Energy-based formulation

In plant cell homeostasis as in many multi-physics systems, using an energy function is a good way to combine the effects of various physical phenomena. Once the complexity of the system physics is embedded in the energy function expression, the resulting system has a clear structure, and the role of each physical effect or transporter is identified. This clear structure is also reflected when it comes to implementing this variational formulation on a computer. In particular, using modern automatic differentiation software, all derivatives of the energy function are automatically evaluated under the hood.

The energy approach makes a distinction between model-specific and generic features, which both contribute to the system evolution. Transport rules, defined by the configuration of transporters, are specific to the proposed guard cell model, but the energy function is more generic. As the sensitivity analysis is based on the energy function only, a similar hierarchy between forces can be expected in other plant cell models. In contrast, formulations using directly ordinary differential equations (Hills et al., 2012; Gerber et al., 2016; Li et al., 2024) do not explicitly separate these two contributions.

### Interpretability of the modeled system

One of the main aims of our modelling strategy was to promote the mechanistic interpretability of the model, in order to obtain insight into the functioning of biological systems. We chose to build a system that acts as a mathematical function, which takes the progress of active transporters as input variables, and returns an output equilibrium state. Thus, we were able to control the rate of the hydrogen pumps present in each membrane and investigate the effect of each active transporter on the model evolution. Interestingly, in the model the relative activities of the plasma membrane and vacuolar membrane hydrogen pumps were the main parameters defining the relative volumes of the vacuolar and cytosolic compartments in the cell. Further, it was possible to define a coordination strategy between the pumps in both membranes to keep the cytoplasmic volume constant while the turgor pressure increases in the cell, as experimentally observed in guard cells during stomata opening (Mirasole et al., 2023). In summary, the model indicates that the activities of the plasma membrane and vacuolar membrane pumps need coordination to achieve biological processes involving changes of volume of the subcellular compartments.

Following our strategy, we performed a sensitivity analysis of the various forces (electrical, chemical and mechanical) acting in our model, and we ranked these forces based on their influence on the system behavior. This gave relevant insight on the functioning of cells with nested membranes and strong turgor pressure, like plant cells. The ranking shows that hydrogen-related forces overcome, by orders of magnitude, the other chemical, electrical and hydraulic forces. Notably, in the model hydraulic forces are subordinate to all the other forces. The importance of hydrogen-related forces depends on the orders-of-magnitude lower hydrogen amounts in both compartments compared to the other chemical species. Under these conditions, small changes in the hydrogen amounts (as those generated by the hydrogen pumps) have amplified consequences on the amounts of the other species transported in the systems. This analysis explains why a transport network based on hydrogen pumps is an efficient strategy to create a large osmotic and turgor pressure in plant cells. The active transport of hydrogens by the pumps creates a leverage effect on the transport of other ions, enabling the building of a strong osmotic pressure. Interestingly, land plants, which are among the organisms with the largest osmotic pressure difference with the extracellular media, evolved a transport network based on hydrogen pumps and hydrogen-coupled exchangers and symporters. In contrast, animal cells maintain a low osmotic pressure difference with the extracellular media and do not exploit hydrogen-coupled transport networks in the plasma membrane, but potassium- and sodium-coupled networks. It is therefore tempting to speculate that this leverage effect is a reason why land plants evolved a transport network based on hydrogen, which is present in low amounts in the cellular and extracellular media.

### Possible extensions of the model

Several extensions can be brought to the proposed model without modifying its principles, as it was the case when adding the buffering mechanism. In this paper, we explored the connections between hydrogen pump activity and the building of a turgor pressure. We considered a cascade of events starting with hydrogen active transport and ending with cell deformation in one dimension. However, this cascade can be extended at both ends. On the upstream side, mechanisms causing the transport of hydrogen may be included in the model. In that case, hydrogen pumps are not controlled by the user, and adenosine triphosphate (ATP), the fuel of active hydrogen transport, is introduced as a new reactant. On the downstream side, more realistic geometric deformation models could be used, leading to more complex terms for the mechanical energy. For instance, a finite-element model could be plugged into our formalism to simulate the change of shape of the guard cell (Woolfenden et al., 2017). At a larger scale, the energetic approach can be used to simulate the regulation of turgor pressure in mechanical models of tissue growth.

### Quasi-static limit and possible generalization to dynamical model

We chose a quasi-static approach to model the system evolution in a simple way. This approach is convenient for models with few transporters, but other choices should be considered to describe more complex transporter configurations. Strong mathematical hypotheses should indeed be satisfied so that the output equilibrium state does not remain constant whatever the value of input variables. In the particular case of Figure 4, a necessary condition for **n**^∗^(*x*) to change as a function of *x* is that the subspaces *V*_*x*_ and 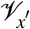 be distinct for two values *x≠ x*′, i.e. the passive transporters do not allow the system to come back to its initial state **n**^0^. In contrast, if the direction **v**_1_ is collinear to **a** in the figure, then **n**^0^ is an element of *V*_*x*_ for every *x* in [0, *x*_max_]. Therefore, the energy minimizer in *V*_*x*_ always remains **n**^∗^(*x*) = **n**^0^, and the active transporters have no effect on the system. In general, we noticed that a sufficient condition for active transporters to induce an evolution of the equilibrium state is the following: at least one active direction **a** is not a linear combination of the passive directions (**v**_1_, **v**_2_, · · ·).

Issues may typically appear when the system involves more transporter than species, with passive transporters allowing to reverse the effect of active transporters. If active transporters fail to change the system equilibrium, i.e. the system behavior is entirely due to transporter dynamics, then a time-dependent model should be considered. A simple way to achieve a dynamic model using the energetic approach is given by Gerber et al. (2016). With our notation, the flux *J* across a passive transporter associated with the direction **v** is proportional to the energy derivative along **v**, i.e. *J* = −*α*∇*G*(**n**)^T^**v**, where *α* is a conductance parameter specific to the considered transporter. Compared with a quasi-static model, a dynamic model may be slightly more difficult to interpret, as its output combines influences from the energy function and transporter conductance. However, it can be used to create more complex behaviors, for instance by considering transporters whose dynamics is affected by voltage or pH.

## 6 Supporting material

### S1 Fig. Hessian matrices at the final state of the simulation

This figure is similar to Figure 9, but the Hessian matrices are evaluated at the final state of the simulation **n**^∗^(**x**_max_). In particular, the analysis of Hessian matrices from Figure 9 still apply to these matrices, as similar differences between coefficient orders of magnitude are present here.

## Footnotes

1 Code available on github.com/gmestdagh/quasistatic-guard-cell-homeostasis

## Notes

https://github.com/gmestdagh/quasistatic-guard-cell-homeostasis

### Competing Interest Statement

The authors have declared no competing interest.

### Summary of Updates

I just changed the address of the corresponding author which appears on the deposit page.

