## Supplementary figures and images for "Multi-physics modeling for ion homeostasis in multi-compartment plant cells using an energy function"

### Supplemental Figure 1

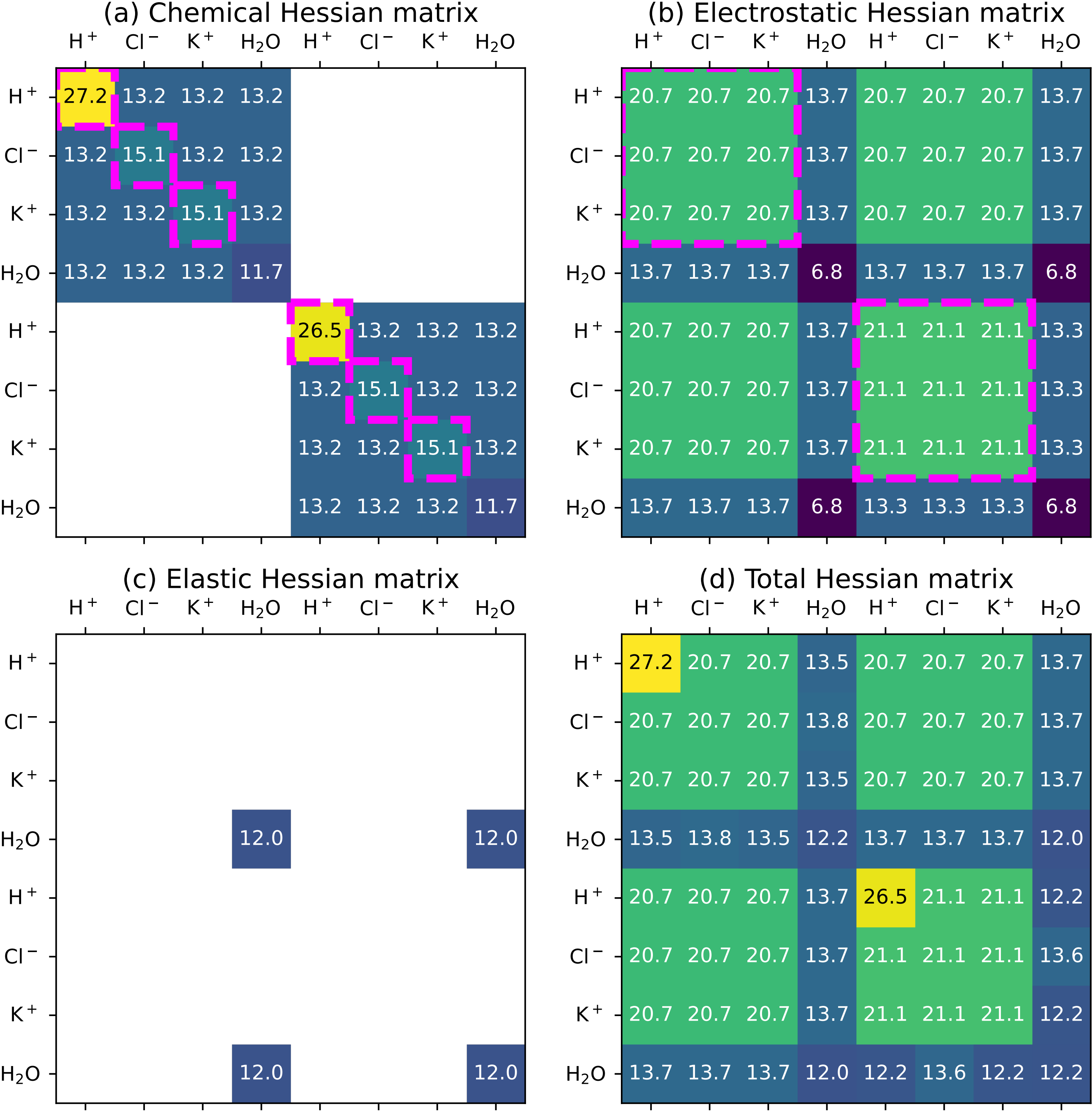
